# Insights into the mechanism of succinimide formation in an archaeal glutaminase

**DOI:** 10.1101/2025.11.17.688757

**Authors:** Anusha Chandrashekarmath, Oishika Jash, Karandeep Singh, Asutosh Bellur, Chitralekha Sen Roy, Aparna Dongre, Nayana Edavan Chathoth, Padmesh Anjukandi, Sanjeev Kumar, Souradip Mukherjee, Padmanabhan Balaram, Sundaram Balasubramanian, Hemalatha Balaram

**Affiliations:** Molecular Biology and Genetics Unit, Jawaharlal Nehru Centre for Advanced Scientific Research, Bengaluru, India; Chemistry and Physics of Materials Unit, Jawaharlal Nehru Centre for Advanced Scientific Research, Bengaluru, India; Department of Chemistry, Indian Institute of Technology, Palakkad-678623, Kerala, India; National Centre for Biological Sciences, Tata Institute of Fundamental Research, Bengaluru, India; Department of Chemistry, Lund University, Sweden; DXBIDT, Lakshmipura, Bengaluru, Karnataka 560073; SCIEX India Private Limited, Hitech defence, Aerospace IT sector, Mahadevakodigehalli village, Bengaluru-562149, India; Advanced Biotech Lab, Ipca laboratories Ltd., Kandivli, Mumbai, India; Department of Chemistry, Washington University in St. Louis, St. Louis, Missouri, USA; Department of Molecular and Cellular Biology, University of Geneva, Geneva, Switzerland

**Keywords:** *Methanocaldococcus jannaschii*, glutamine amidotransferase, succinimide, mass spectrometry, crystal structure, protein thermostability, QM/MM MD simulation

## Abstract

Succinimide (SNN), an intermediate formed during asparaginyl deamidation or aspartyl dehydration in proteins, is generally hydrolysis-prone, leading to isomerization to L/D α/β-aspartyl residue, with the latter being considered deleterious to protein structure and function. An unusually stable SNN-mediated conformational rigidity through restriction of the backbone dihedral angle, ψ, enhances the thermostability of glutamine amidotransferase (GATase) from *Methanocaldococcus jannaschii* (Mj). Although several structural features involved in maintaining a stable SNN and imparting SNN-mediated thermostability have been identified in MjGATase, the residues in the protein that catalyse the rapid and complete conversion of Asn109 to SNN remain unknown. Here, we investigated several site-directed mutants of MjGATase for their ability to retain Asn109 side chain in the unmodified form. Mass spectrometric analysis of 10 single mutants enabled the identification of residues that impacted the proportion of SNN and Asn population in the protein sample. This led to the generation of two double mutants that retained intact Asn109 side chain as observed in the mass spectra and crystal structures. These mutants with intact Asn residue at position 109, displayed lower thermal stability than the protein with the SNN modification. Further understanding of the deprotonation mechanism was addressed using QM/MM MD metadynamics simulations.

**Highlights:**

- Stable succinimide (SNN) arising from deamidation of Asn109 residue imparts hyperthermostability to MjGATase.
- Examination of the structure of MjGATase suggests neighbouring residues playing possible roles in deamidation and cyclization.
- Examination by LC-MS of single site-directed mutants of residues contacting SNN revealed varied levels of intact Asn109 enabling generation of double mutants with complete absence of deamidation.
- Presence of intact Asn109 confirmed by X-ray crystallography highlights the role of Y158, D110, and K151 in mediating SNN formation.
- QM/MM MD metadynamics simulations support experimental findings.

## Introduction

Deamidation, a largely spontaneous post-translational modification (PTM) in proteins occurs at a higher rate in asparaginyl (Asn) compared to glutaminyl (Gln) residues [1–3]. The process of Asn and Gln deamidation leading to Asp/iso-Asp and Glu/iso-Glu residues proceeds through the cyclic intermediates succinimide (SNN) and glutarimide, respectively [3–5]. SNN is also an intermediate during aspartyl dehydration [6,7]. The mechanism of succinimide formation during the deamidation of asparaginyl residues follows three steps namely, (a) deprotonation of the backbone amide NH of the n+1 residue succeeding Asn, (b) nucleophilic attack of the deprotonated amide on the side chain carbonyl carbon of the Asn residue forming a tetrahedral intermediate, and (c) ammonia loss and ring closure leading to SNN formation [8]. The initial step of backbone amide deprotonation is a general-base catalyzed reaction, and the protonation of the leaving ammonia group is a general-acid catalyzed reaction [8]. The cyclic intermediate formed is usually unstable and undergoes hydrolysis to Asp/iso-Asp residues. Racemization of L-SNN to D-SNN followed by hydrolysis can also yield D-Asp/D-iso-Asp [6].

**Scheme 1.**
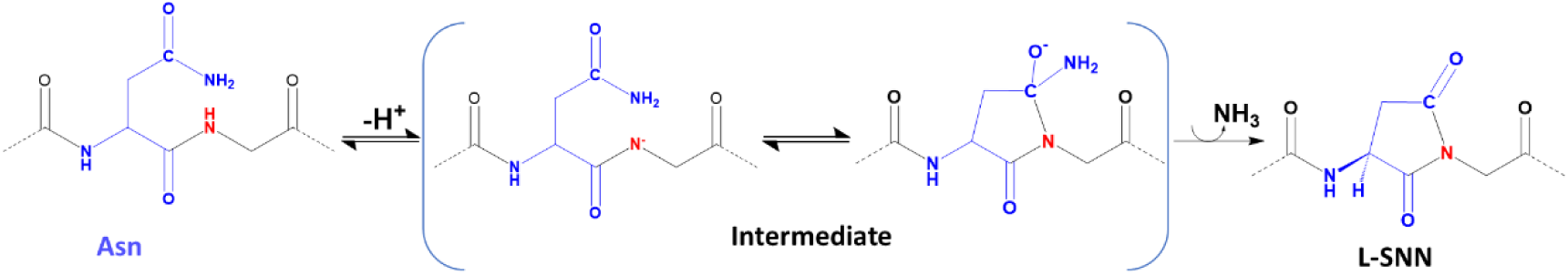
Schematic of succinimide formation by an asparaginyl residue.

The rates of Asn deamidation are well studied in peptides where it has been observed that the rate is influenced by several factors such as temperature, pH, ionic strength, and to a great extent by the nature of the succeeding residue [1,8–11]. Studies done on hexapeptides have shown that the deamidation rate of the Asn residue decreases with an increase in the bulkiness of the side chain of the succeeding residue [6,11]. However, deamidation rates in proteins do not necessarily follow the sequence pattern seen in peptides and are dependent on the three-dimensional structure of the protein [12–15]. The deamidation rates of the Asn residue examined in nearly 120 human proteins showed variation that was governed by both primary sequence and tertiary structures [16]. The effect of three-dimensional structure on deamidation has been studied in a few proteins, such as RNase A [12], trypsin [13], and by MD simulations on the capsid protein of norovirus [17], on triosephosphate isomerase that includes a quantum mechanics/molecular mechanics (QM/MM) free energy calculation providing a mechanistic understanding of the reaction pathway to the tetrahedral intermediate leading to SNN formation [18].

Our earlier studies using mass spectrometry on *Methanocaldococcus jannaschii* glutamine amidotransferase (MjGATase) have shown the role of a remarkably stable SNN, arising from the deamidation of Asn109, in imparting hyper-thermostability to the protein [19]. Mutation of Asn109 to Ser, thereby leading to abrogation of SNN formation, resulted in a protein with reduced thermal stability [19]. A subsequent determination of MjGATase crystal structure enabled the visualization of the SNN modification at residue 109 and its local environment [20]. Structure analysis and enhanced sampling molecular dynamics simulations showed that the SNN induced a conformational lock, and together with long-range interactions contributed to the enhanced thermal stability of the protein. SNN itself was shielded from hydrolysis by the negative charge on the side chain carboxylate of the succeeding D110 residue and, through n-π* interaction with the backbone CO of the preceding residue, E108. These studies provided an insight into the molecular basis of SNN stability to hydrolysis and the role of this PTM in imparting thermal stability to the protein [19,20]. However, the mechanism of the rapid and complete conversion of Asn109 to SNN in MjGATase remains to be explored.

As the three-dimensional structure of the protein plays a major role in deamidation, it is imperative to understand distinct structural features in MjGATase that may promote deamidation. Here, from an analysis of the available structures of MjGATase and its mutants [20], we listed the residues that could be potential internal catalysts mediating SNN formation. These residues directly or indirectly contact the SNN at a 4 Å distance cut-off. Ten site-directed single mutants were generated and were examined for the presence of Asn109 with intact side chain using mass spectrometry (MS). This led to the selection of K151, D110, and Y158, whose mutations in combination may completely abrogate SNN formation. The double mutants D110V_K151L and K151L_Y158F of MjGATase were found by mass spectrometry to largely retain intact the Asn109 side chain and this was validated by structure determination using X-ray crystallography. The MjGATase mutants harboring the intact Asn109 exhibited reduced thermal stability confirming earlier observations [19] on the role of SNN in enhancing the T_m_ values.

The dihedral angles ψ and χ1 of an Asn residue influence succinimide formation [6], demanding a close distance between CG of Asn and the (n+1) backbone N atom [3,21]. Structure based methods have been used to predict deamidation hotspots in antibodies and other proteins [15,22–24]. Using quantities obtained from detailed classical MD simulations such as the solvent accessible surface area (SASA) of the aspartyl side chain, root mean square fluctuations of CA atoms of the aspartyl or asparaginyl residue, and the SASA of (n+1) N and H atoms, and solvent accessibility considering the local environment, the likelihood of degradation mediated through the SNN intermediate has been predicted [25,26]. In addition, the propensity of an asparaginyl residue to undergo deamidation has also been examined through MD simulations combined with quantum mechanical calculations [27]. From the perspective of a mechanistic understanding, there are only a few studies that discuss the deamidation reaction leading to SNN formation from a QM/MM perspective [18,28,29]. Herein, we estimate the activation barrier for the deprotonation step of this PTM based on QM/MM MD simulations through the metadynamics method.

## Results

### Structure analysis suggests residues that may be implicated in the catalysis of SNN formation

The SNN atoms are numbered as shown in Figure 1a. The conversion of an Asn residue to SNN requires a base in the vicinity of the amide NH of the succeeding n+1 residue for proton abstraction. A search for a potential base to abstract the n+1 (D110) amide NH proton was conducted to generate mutants that would be incapable of forming the succinimide and hence retain an intact asparaginyl side chain at residue 109. For this purpose, the contacts of the SNN atoms at a distance cut-off of 4 Å were identified in the structure of wild-type MjGATase (Fig. 1b-d). The MjGATase wild-type (WT) structure will be referred to as MjGATase_SNN_109_, henceforth. A H-bonding interaction was observed between OD1 of SNN109 and L111 NH, in addition to its contacts with D110 OD2, K151 CE, L139 CD1, Y158 CE2, and CG of L111. The SNN109 O contacts E108 carboxy O and K113 CD. The backbone N of D110 which is part of the SNN109, contacts D110 OD1 and OD2, and E108 carboxy O. Of the 4 carbons of the SNN109 ring, CB contacts K107 carboxy O. Although the distance between the amide nitrogen atom (N109, Fig. 1a) of SNN109 and carboxy O of K107 is 3.3 Å indicating an H-bond, the angle criterion for such an interaction does not seem to be satisfied. In addition to these contacts with SNN109, we observed secondary H-bonding interactions between K151 NZ and E137 OE2 and, E137 OE1 and K107 NZ suggesting a possible proton relay.

**Figure 1.**
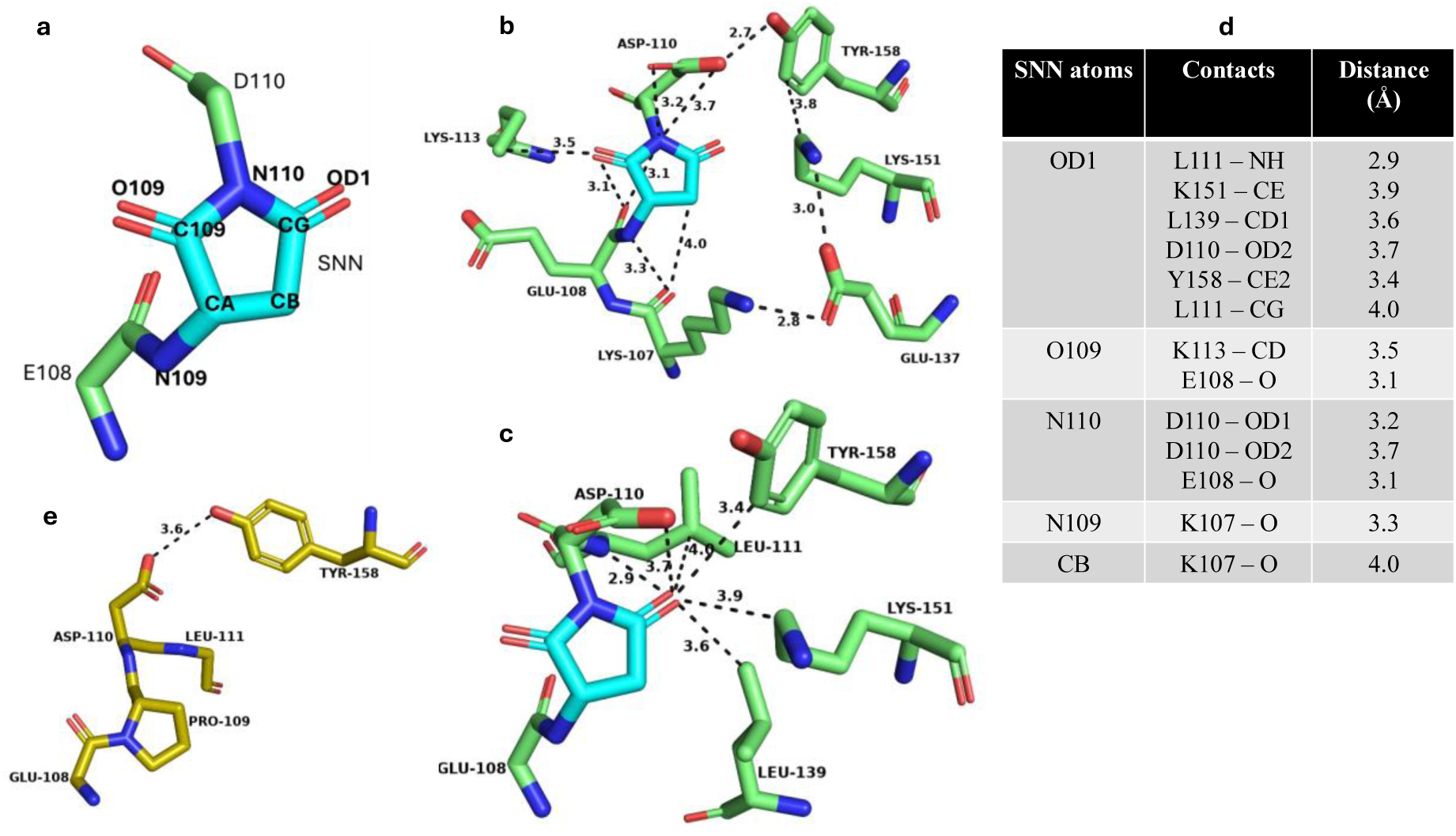
Search for a potential base to abstract the n+1 amide NH proton. (a) Labelling of atoms in SNN. (b), (c) contacts of the succinimide within a 4 Å distance cutoff, in two different views in the MjGATase_SNN109 structure (PDB ID: 7D40). (d) list of contacts made by atoms of SNN with atoms of other residues in MjGATase_SNN109. Contact distances in Å are provided. (e) the contact between Asp110 side chain C=O and Tyr158 OH in MjGATase_N109P structure (PDB ID: 7D97).

The above contact analysis suggested that the residues D110, Y158, K151, E137, K107, E108, and K113 could have a direct or indirect role in the abstraction of the D110 amide proton, initiating SNN formation. Of these residues, as D110 side chain carboxylate interacts with D110 NH (N110, Fig. 1a, b), it appeared as a possible potential base for abstracting the proton from its backbone NH. The D110 residue was mutated to the imino acid, proline, that lacking the amide NH proton would be incapable of forming SNN and, therefore, enable the retention of the Asn109 side chain. D110 was also mutated to Asn and Val to obliterate the negative charge thereby eliminating the basic nature of the side chain.

The second residue identified from the contact analysis that could assist in proton abstraction by functioning as a component of a proton relay is Y158, the side chain OH of which contacts D110 carboxylate (Fig. 1b). Interestingly, even in the mutant, MjGATase_N109P, the H-bond between D110 carboxylate and Y158 side chain OH continues to be retained (Fig. 1e). Similarly, a proton relay could also occur between the residues K151, E137 and K107 (Fig. 1b), hence, the site directed mutants Y158F, K151L, E137L, and K107L were generated.

Although a direct contact between the side chain of E108 and SNN is not observed, it was mutated to leucine and glutamine as E108 precedes SNN and could have implications in both the formation and stability of this PTM. The last residue identified from the contact analysis is K113, whose side chain CD is within the 4 Å distance cut-off from SNN (Fig. 1b) and this was mutated to Ala. However, it should be noted that the electron density of the entire side chain of K113 is absent in MjGATase_SNN_109_ (PDB ID: 7D40), MjGATase_N109P (PDB ID: 7D96), and MjGATase_D110G (PDB ID: 7D95) structures.

### MS analysis of single site-directed mutants indicates possible roles for D110, Y158, K151, K113, and E108 in SNN formation

The single mutants of MjGATase E108L, E108Q, K107L, E137L, K151L, K113A, Y158F, D110N, D110P, and D110V were generated and confirmed by DNA sequencing. The MjGATase_SNN_109_ and mutant proteins were purified to homogeneity and subjected to intact protein mass analysis. Figure 2, Figure S1, and Figure S2a show intact protein mass spectra of the MjGATase_SNN_109_ and all the mutant proteins. As MjGATase_SNN_109_ has a stable SNN at position 109, we expect the following protein populations to be seen in the mass spectra of the MjGATase_SNN_109_ and mutants, *viz*., population with SNN having a mass that is 17 Da lower than the expected mass (M-17), population with intact Asn having the expected mass (M), and SNN hydrolyzed species having a mass that is 1 Da higher than expected mass (M+1).

**Figure 2.**
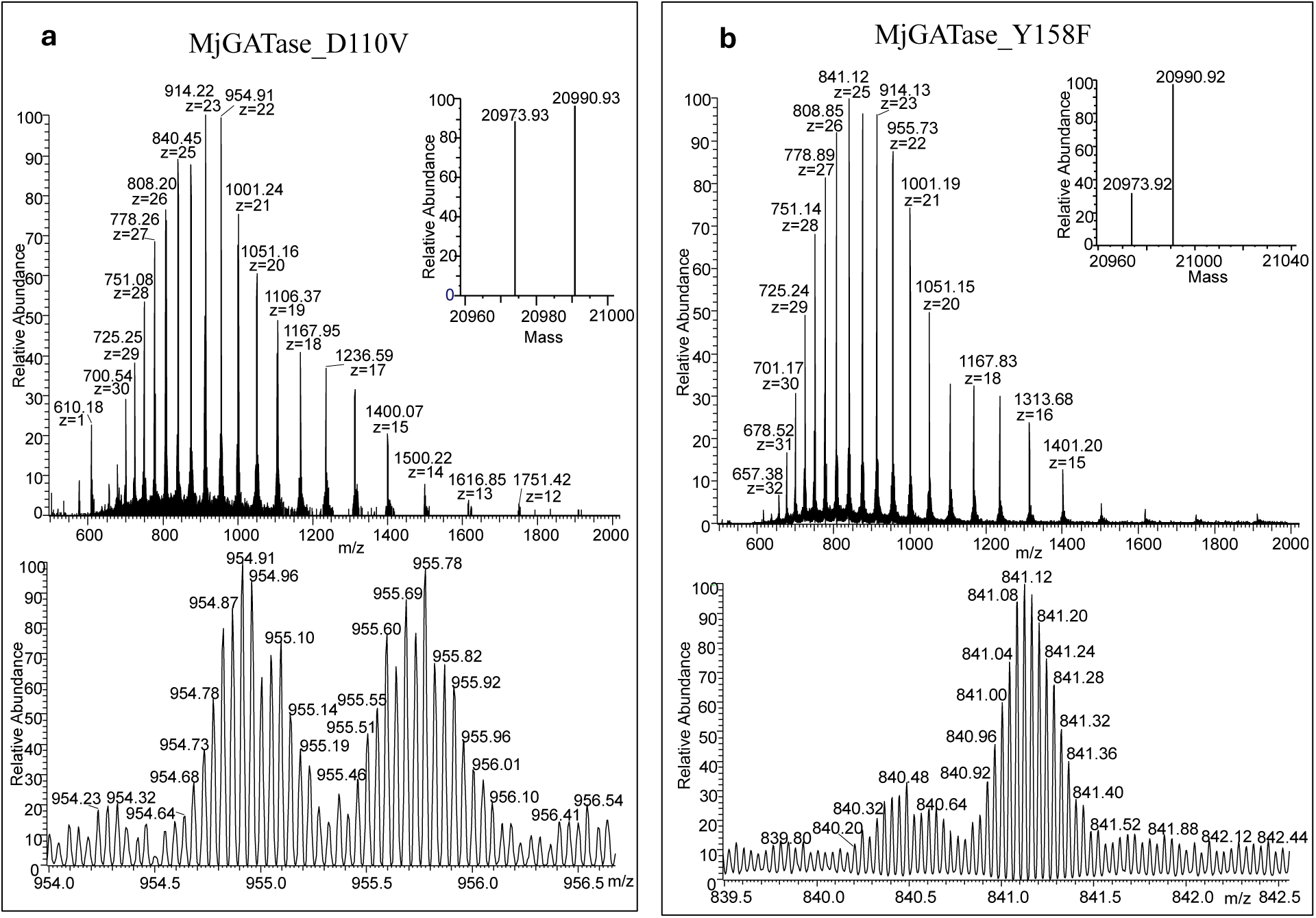
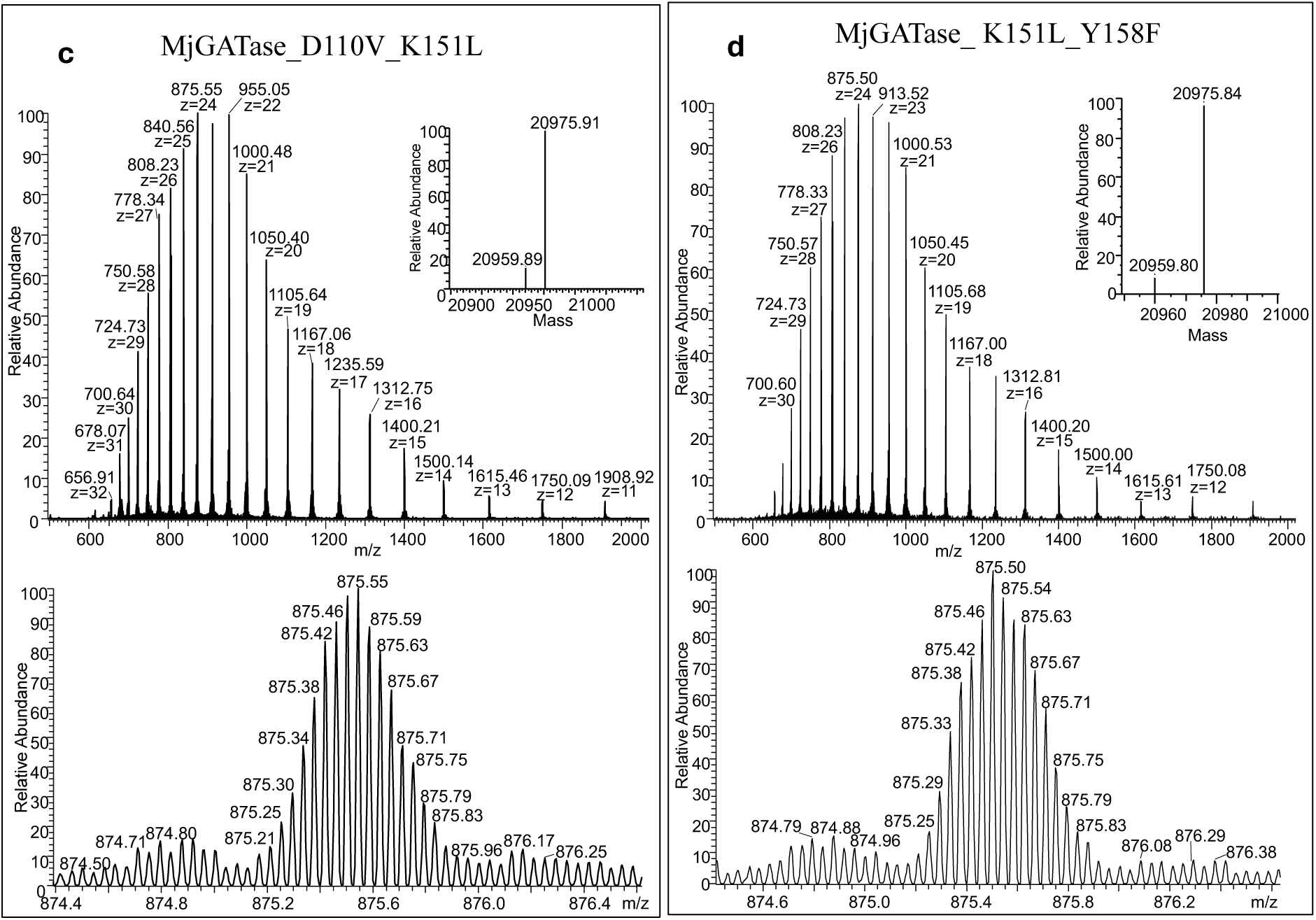
Mass spectra of mutants of MjGATase that have retained the side chain of Asn109. In each main panel, the top sub panel shows the entire mass spectrum of the protein with the inset showing the monoisotopic mass obtained after deconvolution. The bottom panel shows the expanded spectrum of a single charge state. (a) LC-MS of MjGATase_D110V (Mcalc 20990.92 Da). Deconvoluted spectrum in the inset shows equal populations with Mobs value of 20973.93 Da corresponding to SNN109 and Mobs of 20990.93 Da corresponding to the expected mass with Asn109. (b) LC-MS of MjGATase_Y158F (Mcalc 20990.89 Da). Deconvoluted spectrum in the inset shows a major population having a Mobs value of 20990.92 Da corresponding to the expected mass with Asn109 and a small population of 20973.92 Da corresponding to SNN109. (c) LC-MS of MjGATase_D110V_K151L (Mcalc 20975.91 Da). Deconvoluted spectrum in the inset shows a major population with Mobs value of 20975.91 Da corresponding to the expected mass with Asn109 and an unassigned minor population with Mobs of 20959.89 Da which is 16 Da lower than the expected mass of the protein. (d) LC-MS of MjGATase_K151L_Y158F (Mcalc 20975.88 Da). Deconvoluted spectrum in the inset shows a major population having Mobs value of 20975.84 Da corresponding to the expected mass with Asn109 and an unassigned minor population having Mobs value of 20959.80 Da which is 16 Da lower than the expected mass of the protein.

#Table S1 summarizes the results obtained from intact protein mass spectrometric analysis of MjGATase_SNN_109_ and mutant proteins. The various species of MjGATase mutants observed were those having SNN, intact Asn, SNN hydrolyzed to D/isoD, and methionine oxidised species. The relative abundance of these species was obtained from the MS data with an abundance cut-off set between 10-15% above the noise level. As expected, MjGATase_SNN_109_ showed a major population corresponding to the presence of SNN (Fig. S1a) while MjGATase_E108L, MjGATase_E137L, MjGATase_K107L and MjGATase_K151L (Fig. S1b-e) also showed a major SNN containing population. The first mutant to show a population of protein with intact Asn, albeit minor, was MjGATase_K151L. MjGATase_K113A (Fig. S1g) also showed a minor population with Asn intact with the rest corresponding to protein with SNN. With a predominant SNN population, MjGATase_E108Q (Fig. S1h) also behaved similar to MjGATase_E108L but retained a small population of intact Asn. MjGATase_D110V showed two populations in almost equal proportions corresponding to Asn109 intact and SNN, respectively (Fig. 2a). A dramatic increase in intact Asn population was seen in MjGATase_Y158F (Fig. 2b) where the SNN population was 31.7% lower in relative abundance than the non-cyclized Asn109 protein (Table S1). MjGATase_D110N showed a major population corresponding to D/isoD (Fig. S1f). As expected, ESI-MS of MjGATase_D110P with an imino acid as the n+1 residue showed complete retention of only the intact Asn109 population (Fig. S2a). To summarize, the intact Asn109 containing population was the highest in Y158F, followed by D110V that had equal amounts of Asn109 and SNN, and finally a small proportion of Asn109 in K151L, K113A, and E108Q mutants of MjGATase, hinting at the role of these residues in SNN formation.

### Confirmation by MS/MS analysis of peptides containing SNN109/Asn109

Due to the questionable reliability in precisely differentiating a mass difference of 1 Da between Asn intact (M Da) and hydrolyzed SNN (M+1 Da), the assigned Asn-intact population observed in the MjGATase mutants K151L, K113A, Y158F, and D110V/D110N could be due, at least partially, to D/iso-D population or vice-versa. Therefore, to determine the true Asn109 intact population, in-gel trypsin digestion followed by MS/MS analysis of the derived peptides was carried out. The data are summarized in Table S2. In all the five mutants, the VYVDKEN_109_D/V/N_110_LFK/A_113_NVPR peptide was found to exist as Asn109, SNN109, or D/isoD109, albeit in varying amounts (Fig. 3a and b, Fig. S3a-c, Fig. S4a-f, Fig. S5a-d). The relative quantification of different populations of a peptide was made using the number of peptide-spectrum matches (PSM) [30,31], and peptides with at least 2 PSMs and a maximum of 2 missed cleavages were included in the analysis. In MjGATase_K151L, the 109^th^ residue with intact Asn was low at 5%, whereas in K113A the percentage of intact Asn109 increased to 13% (Table S2). The MjGATase_D110N also had an Asn109 intact population of 19% (Table S2). In D110V and Y158F mutants, the percentage further increased to 27 and 45%, respectively (Table S2). In comparison, in MjGATase_SNN_109_, the percentage of intact Asn109 was below 5% (Fig. S3d, Fig. S5e and f). As expected from the D110P mutant, all the peptides containing the 109^th^ residue had only Asn109 (Fig. S2b). In summary, mutation of the residues D110 and Y158 significantly impacts the levels of the intact Asn109 population. Examination of the MjGATase_SNN_109_ structure shows the possibility of a proton relay between side chains of Y158 and K151 (Fig. 1b). Hence, the two double mutants D110V_K151L and K151L_Y158F were generated to examine the levels of intact Asn109 population.

**Figure 3.**
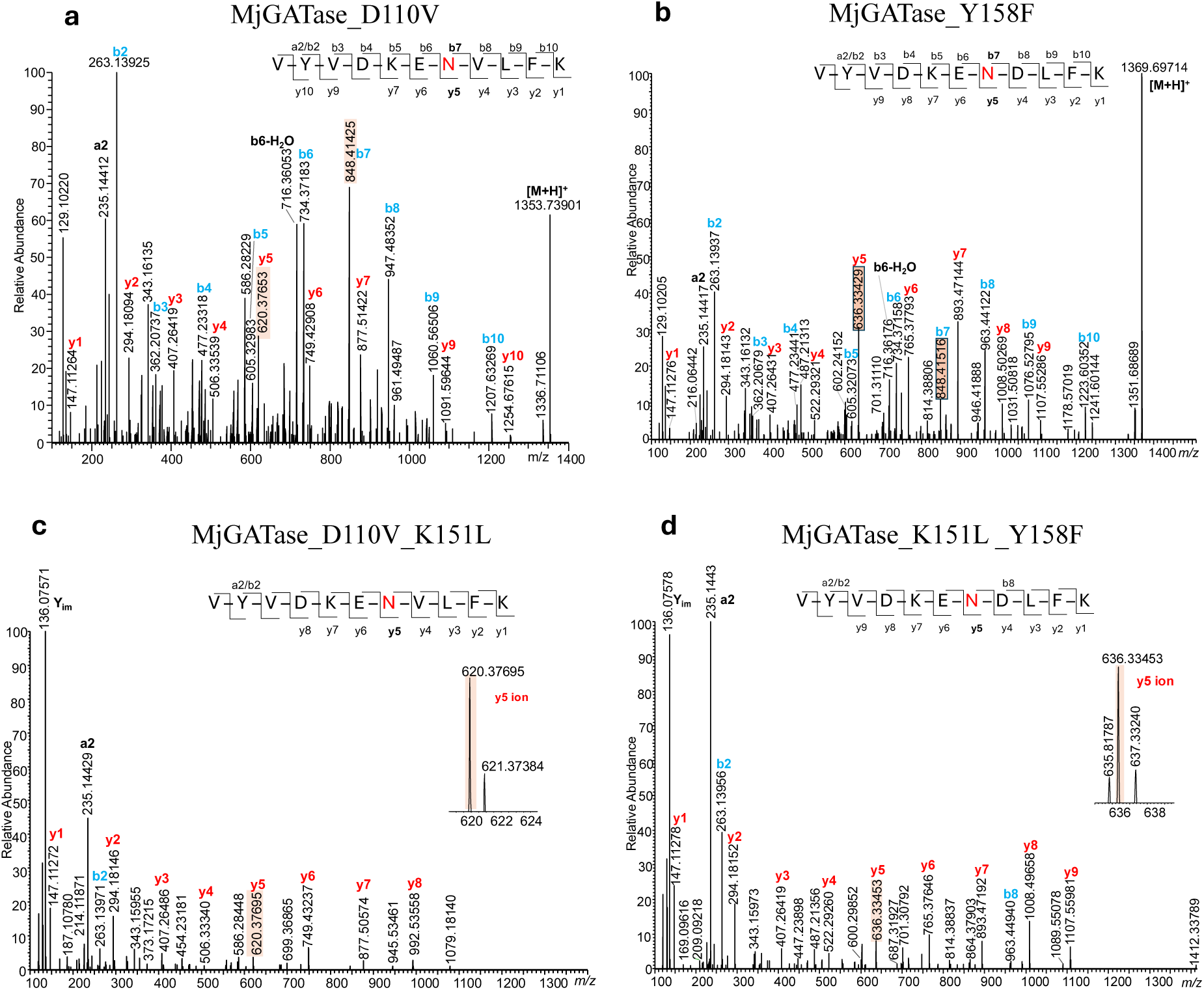
HCD-MS/MS analysis of in-gel trypsin digested peptides show Asn109 intact in specific mutants. (a) MS/MS of peptide of m/z 1353.73 derived from tryptic digest of MjGATase_D110V shows the presence of intact Asn109. The y5 and b7 ions with m/z values of 620.37 and 848.41, respectively are highlighted confirming Asn109. The y4 and b8 ions with m/z values of 506.33 and 947.48, respectively also confirm the mutation of Asp to Val at residue 110. (b) MS/MS of peptide of m/z 1369.69 derived from tryptic digest of MjGATase_Y158F shows the presence of intact Asn109. The y5 and b7 ions with m/z values of 636.33 and 848.41, respectively are highlighted confirming Asn109. (c) MS/MS of triply charged peptide of m/z 451.918 derived from tryptic digest of MjGATase_D110V_K151L shows the presence of intact Asn109. The y5 ion of Asn109 with m/z 620.37 is highlighted and the inset shows the isotope distribution of this ion. The y4 ion with m/z 506.33 also confirms the mutation of Asp to Val at residue 110. (d) MS/MS of doubly charged peptide of m/z 685.35 derived from tryptic digest of MjGATase_K151L_Y158F shows the presence of intact Asn109. The y5 ion of Asn109 with m/z 636.33 is highlighted and the inset shows the isotope distribution of the y5 ion. MS/MS spectra of corresponding peptides with SNN109 or D/isoD109 are provided in Fig. S4a-d for MjGATase_D110V and MjGATase_Y158F and Fig. S6 for the two double mutants.

### MS and MS/MS analyses of double mutants of MjGATase show high levels of intact Asn109

The intact protein mass spectra of D110V_K151L and K151L_Y158F mutants of MjGATase showed a high population of intact Asn109 (Fig. 2c and d, Table S1). To further confirm that the mass corresponds to Asn109 intact and not D/isoD109, in-gel trypsin digestion followed by MS/MS analysis of the peptides was carried out. Although SNN109 or D/isoD109 populations of the peptide VYVDKEN_109_D/V_110_LFK were observed (Fig. S6), their total amounts corresponded to 12.6% for D110V_K151L mutant and 18% for K151L_Y158F mutant with the Asn109 being the major population (Table S2). In MjGATase_D110V_K151L, the 109^th^ residue with Asn contributed to 87.4%, whereas in K151L_Y158F mutant, the percentage was 82% (Table S2). The MS/MS analysis reassured that in the double mutants with Asn109 intact (Fig. 3c and d), deamidation and SNN formation have not occurred to a significant extent. In addition, the agreement seen across the intact protein MS analysis and peptide MS/MS analysis added further confidence to the findings. It must be noted, that as the masses of the two double mutants are identical, the protein samples were subjected to in-gel tryptic digestion and the presence of the peptides carrying the expected mutations was confirmed (Fig. S7).

### Crystal structures of D110P, D110V_K151L, and K151L_Y158F mutants of MjGATase confirm the presence of intact Asn109

As mass spectrometric data of MjGATase_D110P, MjGATase_D110V_K151L, and MjGATase_K151L_Y158F showed a complete or largely complete homogenous population with Asn109 intact, these were chosen for structure determination by X-ray crystallography. The crystals of MjGATase_D110P, MjGATase_D110V_K151L, and MjGATase_K151L_Y158F diffracted to 2.4, 2.5, and 2.4 Å resolution with the space group P3121, P41, and P41, respectively, and the structures were determined by molecular replacement using MjGATase_SNN_109_ structure as the model (PDB ID: 7D40). The data collection and refinement statistics are provided in Table 1.

**Table 1.**
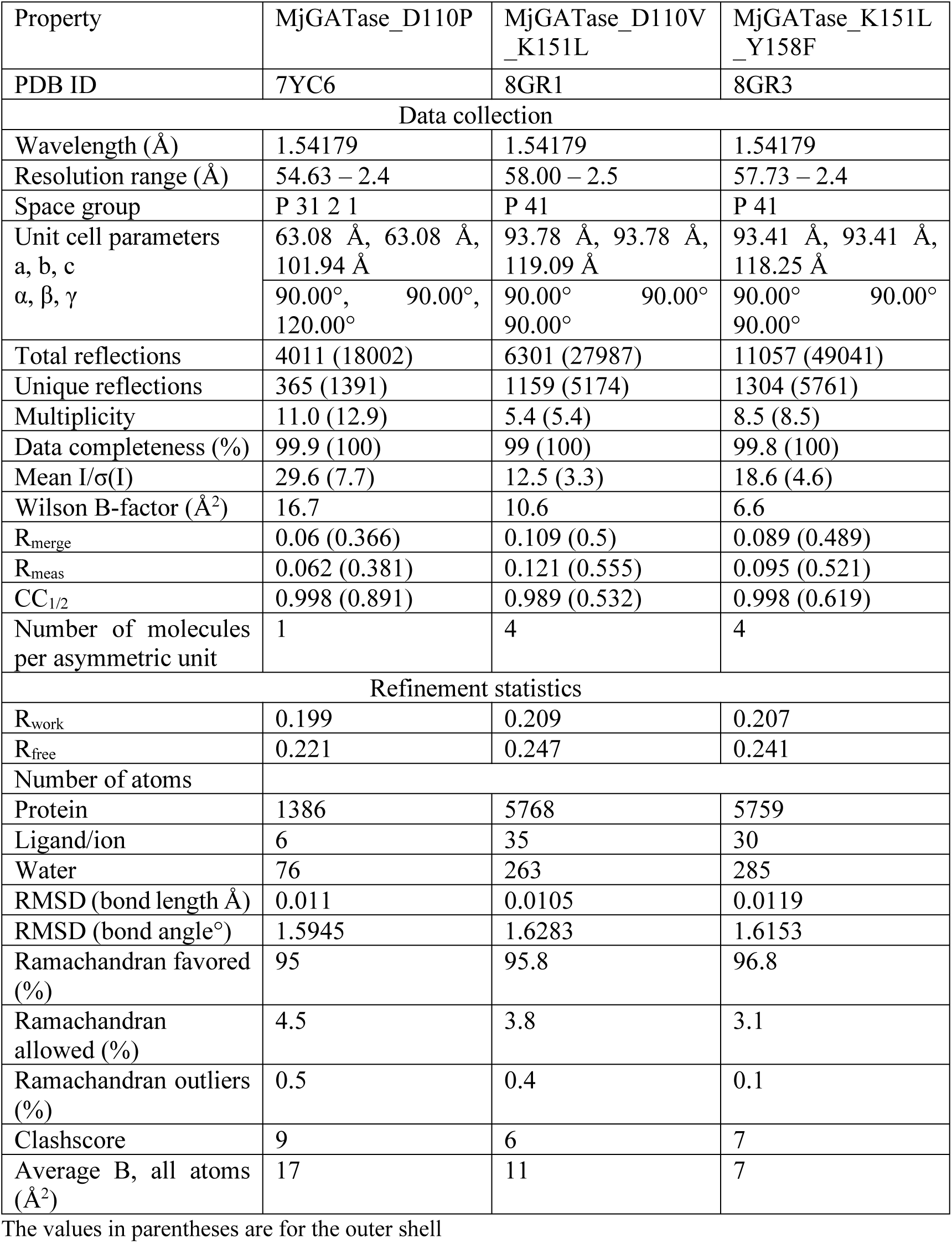
Summary of data collection and refinement statistics.

| Property | MjGATase_D110P | MjGATase_D110V_K151L | MjGATase_K151L_Y158F |
| --- | --- | --- | --- |
| PDB ID | 7YC6 | 8GR1 | 8GR3 |
| Data collection |  |  |  |
| Wavelength (Å) | 1.54179 | 1.54179 | 1.54179 |
| Resolution range (Å) | 54.63 – 2.4 | 58.00 – 2.5 | 57.73 – 2.4 |
| Space group | P 31 2 1 | P 41 | P 41 |
| Unit cell parameters<br>a, b, c<br>$\alpha$, $\beta$, $\gamma$ | 63.08 Å, 63.08 Å,<br>101.94 Å<br>90.00°, 90.00°,<br>120.00° | 93.78 Å, 93.78 Å,<br>119.09 Å<br>90.00° 90.00°<br>90.00° | 93.41 Å, 93.41 Å,<br>118.25 Å<br>90.00° 90.00°<br>90.00° |
| Total reflections | 4011 (18002) | 6301 (27987) | 11057 (49041) |
| Unique reflections | 365 (1391) | 1159 (5174) | 1304 (5761) |
| Multiplicity | 11.0 (12.9) | 5.4 (5.4) | 8.5 (8.5) |
| Data completeness (%) | 99.9 (100) | 99 (100) | 99.8 (100) |
| Mean I/$\sigma$(I) | 29.6 (7.7) | 12.5 (3.3) | 18.6 (4.6) |
| Wilson B-factor (Å<sup>2</sup>) | 16.7 | 10.6 | 6.6 |
| R<sub>merge</sub> | 0.06 (0.366) | 0.109 (0.5) | 0.089 (0.489) |
| R<sub>meas</sub> | 0.062 (0.381) | 0.121 (0.555) | 0.095 (0.521) |
| CC<sub>1/2</sub> | 0.998 (0.891) | 0.989 (0.532) | 0.998 (0.619) |
| Number of molecules<br>per asymmetric unit | 1 | 4 | 4 |
| Refinement statistics |  |  |  |
| R<sub>work</sub> | 0.199 | 0.209 | 0.207 |
| R<sub>free</sub> | 0.221 | 0.247 | 0.241 |
| Number of atoms |  |  |  |
| Protein | 1386 | 5768 | 5759 |
| Ligand/ion | 6 | 35 | 30 |
| Water | 76 | 263 | 285 |
| RMSD (bond length Å) | 0.011 | 0.0105 | 0.0119 |
| RMSD (bond angle°) | 1.5945 | 1.6283 | 1.6153 |
| Ramachandran favored<br>(%) | 95 | 95.8 | 96.8 |
| Ramachandran<br>allowed (%) | 4.5 | 3.8 | 3.1 |
| Ramachandran outliers<br>(%) | 0.5 | 0.4 | 0.1 |
| Clashscore | 9 | 6 | 7 |
| Average B, all atoms<br>(Å<sup>2</sup>) | 17 | 11 | 7 |

The values in parentheses are for the outer shell

In contrast to MjGATase_SNN_109_, in the structures of the three mutants, the electron density for Asn109 was clearly evident (Fig. 4 and S2c), corroborating the findings from mass spectrometry. The overall structures of the mutants superpose very well on the MjGATase_SNN_109_ structure with root mean square deviation (RMSD) of 0.412 Å for MjGATase_D110P, 0.3 Å for MjGATase_D110V_K151L and 0.24 Å for MjGATase_K151L_Y158F. The superposition of all the three mutant structures on the WT is shown in Figure 5a.

**Figure 4.**
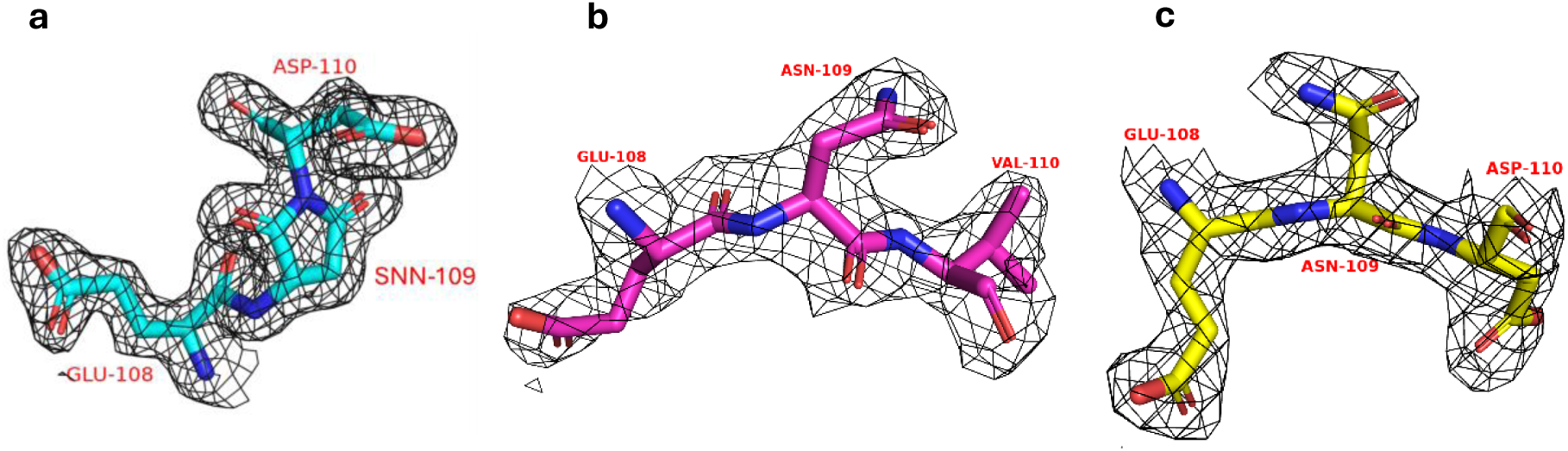
Structural evidence for the presence of intact Asn109 side chain in the site-directed mutants. The 2Fo-Fc electron density map contoured to 1σ (black mesh) for residues 108, 109, and 110 in the structures of (a) MjGATase_SNN109, (b) MjGATase_D110V_K151L, and (c) MjGATase_K151L_Y158F. The electron density is evident for Asn109 in the structures of the mutants.

**Figure 5.**
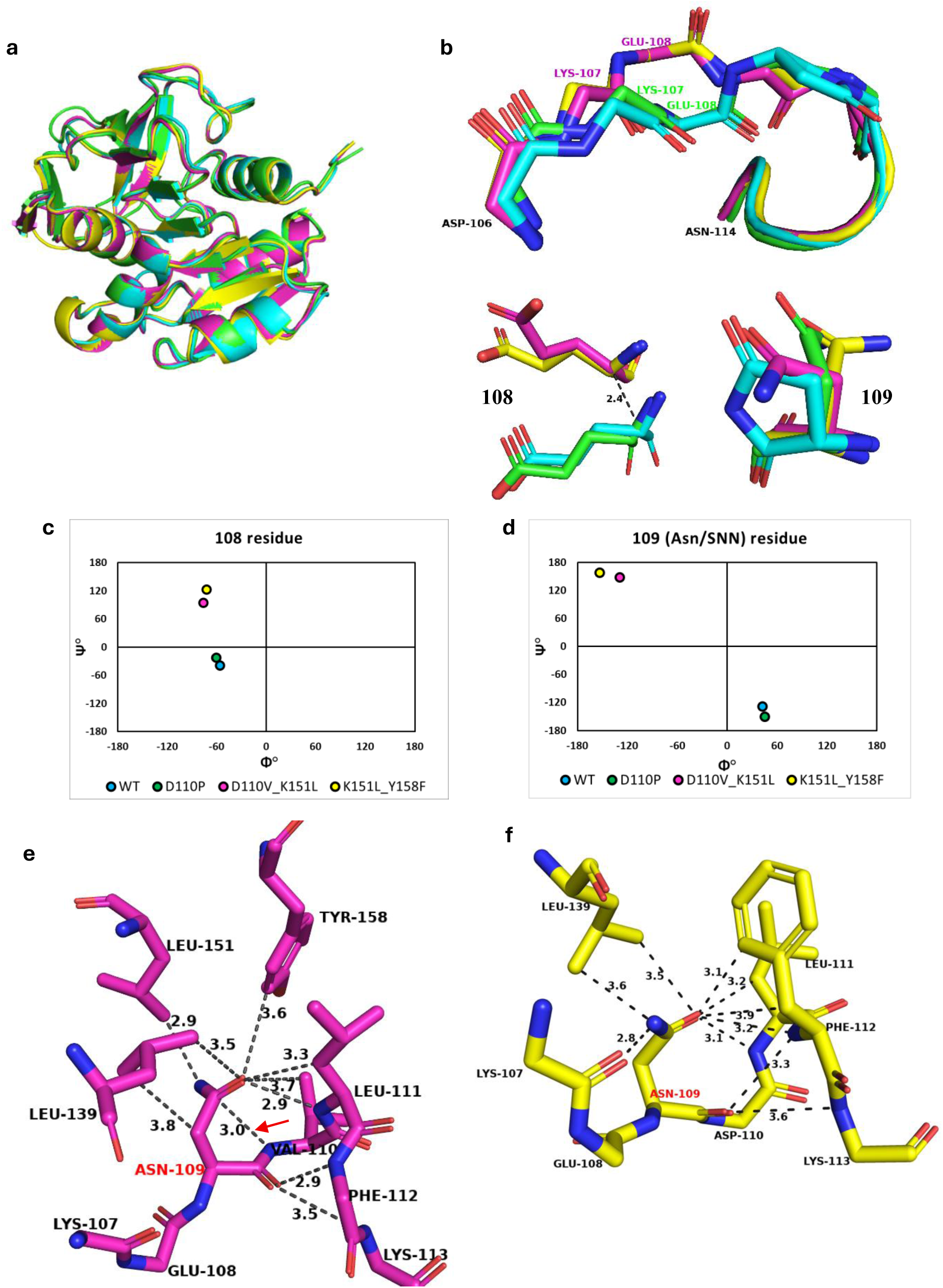
Structural analysis of MjGATase_SNN109 and mutants. (a) Superposition of the structures of the mutants D110P (green), D110V_K151L (purple), and K151L_Y158F (yellow) on the MjGATase_SNN109 (cyan) did not show dramatic differences in the overall structure. (b) The top panel shows the superposition of the loop harbouring Asn109/SNN109 from residues 106 to 114. The backbone of residues from 106 to 110 are shown in sticks and residues 111 to 114 are shown in cartoon representation. The bottom panels show the zoomed in view of the superposition of the residues 108 and 109 along with their side chains. The non-superimposability of the CA of E108 in the MjGATase_SNN109 on to their counterpart in the double mutants shows the lateral movement of this segment in the structures of the mutants. The dashed line indicates the movement of E108 by 2.4 Å. Note that E108 in the structure of MjGATase_D110P overlays well on to E108 in the MjGATase_SNN109 structure. Colours used are as indicated in panel (a). (c) Ramachandran map occupancy of the residue 108 in WT and mutants. (d) Ramachandran map occupancy of residue 109 in WT and mutants. WT in (c) and (d) refers to MjGATase_SNN109 structure. Contacts of Asn109 at 4 Å distance cut-off in the structures of (e) MjGATase_D110V_K151L and (f) MjGATase_K151L_Y158F. The arrow in (e) points to the near-attack conformation distance between the backbone amide of Val110 and CG of Asn109.

### Structure analysis of the mutants

The loop (106-114 residues) harboring Asn109 in MjGATase_D110P superposes well with that in the MjGATase_SNN_109_ structure. However, in the structure of the double mutants, there is a variation in the backbone conformation from residues K107 to L111 (Fig. 5b). This shift is clearly evident in the dihedral angles, φ and ψ for the residues E108 and Asn109 in the structures of the double mutants MjGATase_D110V_K151L and MjGATase_K151L_Y158F, wherein they occupy different quadrants of the Ramachandran map compared to that in the MjGATase_SNN_109_ structure (Fig. 5c and d). In the WT structure, the dihedral angles φ (42.6°) and ψ (-128.8°) of SNN109 fall in the bottom right quadrant of the Ramachandran map. The MjGATase_D110P structure has the Asn residue taking up φ and ψ values of 45.1° and -150.5°, respectively lying in the bottom right quadrant of the Ramachandran map as seen for the WT enzyme. In contrast, in MjGATase_D110V_K151L and MjGATase_K151L_Y158F structures, the dihedral angles of the corresponding Asn109 fall in the top left quadrant of the Ramachandran map with values being φ -129.3°, ψ 148.6°, and φ -153.6°, ψ 157.8°, respectively (Fig. 5d).

The SNN-containing loop in the MjGATase_SNN_109_ has an α-turn stabilized by a hydrogen bond between Glu108 CO-Phe112 NH and a β-turn with a hydrogen bond between Glu108 CO-Leu111 NH (Fig. S8a). These features are retained in the D110P mutant (Fig. S2d). This α-turn and β-turn initiating from E108 in the WT is replaced by an α-turn and β-turn initiating from Asn109 in D110V_K151L and K151L_Y158F MjGATase mutants. The α-turn involves Asn109, Val110/Asp110, Leu111, Phe112 and Lys113 with hydrogen bonding interaction between Asn109 CO-Lys113 NH and β-turn involving Asn109, Val110/Asp110, Leu111 and Phe112 with hydrogen bonding interaction between Asn109 CO-Phe112 NH (Fig. S8b and c). The α-turn in the MjGATase_SNN_109_ is facilitated as E108 backbone CO points inwards into the loop with φ = -56.3°, ψ = -38.7°, whereas in the double mutants, a change in the dihedral angle, ψ for the residue E108 to 94.2° and 122.5° in MjGATase_D110V_K151L and MjGATase_K151L_Y158F respectively, leads to a reorientation of backbone CO (Fig. 5c and S8). The conformation adopted by Asn109 being dissimilar to that of the SNN109, has altered the backbone orientation of the preceding residue, E108 in the double mutants (Fig. 5b).

In the three structures where Asn109 is intact, the Asn109 side chain was found in three different conformations (Fig. 5b, bottom right panel). The side chain dihedral angles χ1 and χ2 of Asn109 are 72.9° and 130.8°, 173.9° and –99.6°, 58.2° and 93.1° in MjGATase_D110P, MjGATase_D110V_K151L and MjGATase_K151L_Y158F, respectively. The corresponding dihedral angles χ1 and χ2 in the WT harboring SNN are 127.3° (χ1) and 179.4° (χ2). Superposition of the loop harboring SNN/Asn in WT and the mutant structures showed that Asn109 conformation in K151L_Y158F MjGATase was drastically different from that of SNN whereas, in the D110V_K151L MjGATase structure, it was closest to the SNN conformation mimicking the ‘near-attack’ conformation (χ1 of +120° and χ2 of +90° or -90°, [3]) (Fig. 5b, bottom right panel). The distance proposed between the backbone amide of the n+1 residue and CG of Asn (n^th^ residue) for the ‘near-attack’ conformation favoring cyclization is < 3.5 Å [3,18,32,33] and the corresponding distance in the D110V_K151L mutant of MjGATase structure is 3.0 Å (Fig. 5e).

As shown in the MjGATase_SNN_109_ structure, most of the contacts arising from SNN involve the OD1 atom (Fig. 1c and d). Similarly, in the double mutant structures, most of the contacts of Asn109 side chain were with the OD1 atom (Fig. 5e and f). In both the double mutant structures, OD1 of Asn109 makes a H-bonding interaction with the backbone NH of L111 residue (Fig. 5e and f). This contact is retained after SNN formation as observed in the MjGATase_SNN_109_ structure and retained in the D110P structure where Asn109 cannot form SNN (Fig. 1c and S2e). In the mutant K151L_Y158F, the OD1 atom of Asn109 makes an additional H-bonding interaction with backbone NH of F112 and ND2 makes H-bonding interaction with backbone CO of K107 residue (Fig. 5f). These additional H-bonds of Asn109 in the K151L_Y158F structure could be the reason for the Asn109 conformation being different from the Asn109 conformation in the D110V_K151L structure. The other interactions of the OD1 atom in both the double mutant structures are van der Waals in nature.

The acidity of backbone amide hydrogen of the n+1 residue is mainly dictated by the backbone dihedral ψ adopted by the n+1 residue [18,34]. The dihedral angles adopted by V110 are φ = -46.3°, ψ = -34.1° in the structure of MjGATase_D110V_K151L and φ = -59.3°, ψ = - 32.8° for residue D110 in the structure of MjGATase_K151L_Y158F. The corresponding backbone dihedrals for D110 are φ = -100.2°, ψ = -47.5° in the MjGATase_SNN_109_ structure. In the classical MD run of MjGATase_ASN_109_ (discussed below), the ψ value was in the range of -10° to -50° for D110 (Fig. S9a). In the plot of relative proton affinity (2-acetamido-N-methylacetamide and N-formyl-glycinamide) as a function of dihedral angles [18,34], these conformations lie in moderately low proton affinity regions, hence, the backbone amide hydrogen of the n+1 residue is moderately acidic in MjGATase.

### Mutants with intact Asn109 exhibit lowered thermal stability

The absence of the SNN resulted in lower melting temperatures for mutants MjGATase_D110P, MjGATase_D110V_K151L, and MjGATase_K151L_Y158F with T_m_ values ranging between 82-84 ℃ for the mutants and >100 ℃ for WT [19] as shown in Figure 6a. The rate of unfolding at 80 ℃ was also measured for the double mutants and the WT protein wherein both double mutants within a few minutes showed rapid unfolding with a sharp decline in the slope of the unfolding curve whereas the WT protein did not unfold even after 180 minutes (Fig. 6b). REST2 MD simulations on mutants where Asn109 was replaced by Ser and Pro showed structural destabilization, at higher temperatures, of the distal β-sheet originating from the SNN loop [20]. Taken together, the current study also implicates a critical role for the SNN in maintaining hyperthermostability.

**Figure 6.**
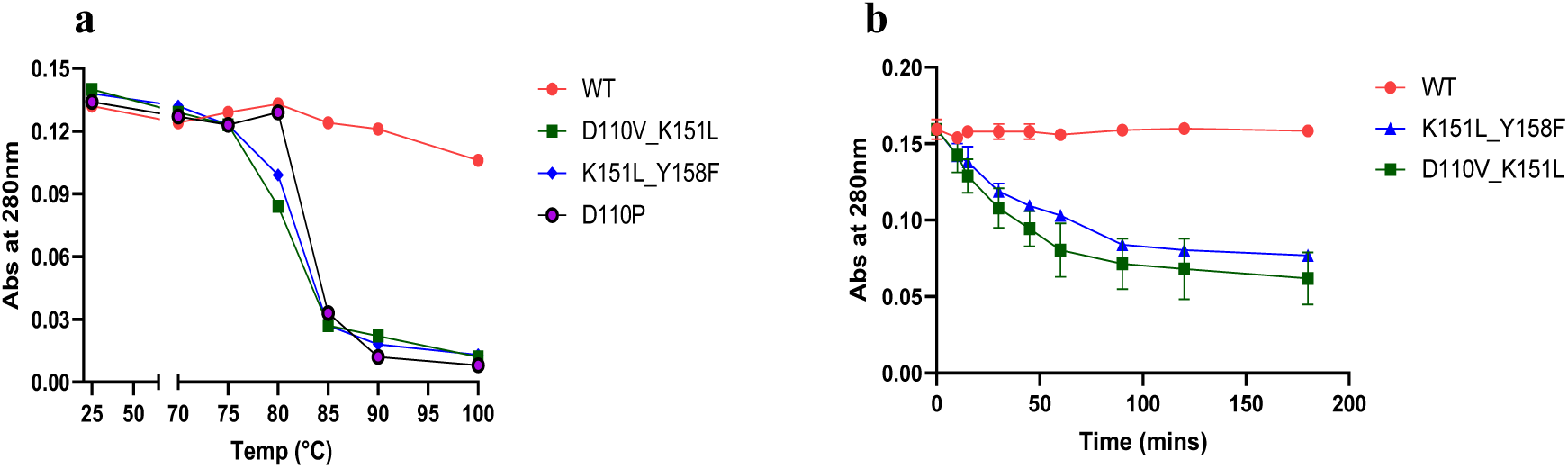
Modification of Asn109 to an SNN is required for the hyperthermostability of MjGATase. Comparison of the stabilities of MjGATase_SNN109 (WT) and the mutants D110P, D110V_K151L, and K151L_Y158F. (a) Thermal melting curve of the mutants and WT. The experiment was repeated twice, and similar Tm values were obtained in both repeats. One representative graph is shown in the figure. Temp along X-axis denotes temperature. (b) The rate of unfolding plotted for the double mutants and WT, the values are mean with error bars as SEM (n=2).

### Formation of SNN at low pH in MjGATase_ 110V_K151L

To investigate if the intact Asn109 population of the mutant D110V_K151L can convert to SNN at lower pH, the WT and the mutant were incubated at pH 3 for 16.5 hours at 60 ℃ and examined using mass spectrometry. The mass spectrum of MjGATase_D110V_K151L showed an increase in the SNN109 population from 17 to 34% over a period of 16.5 hours relative to the Asn109 containing population (Fig. S10a and b). The presence of the SNN population shows the ability of the mutant to form SNN at low pH, albeit at a very slow rate. This detection of SNN despite the absence of the putative catalytic base (D110) in the mutant indicates that the overall active site geometry is optimized for very high rates of deamidation, hence the minor increase in SNN levels over prolonged incubation at low pH. It should be noted that under crystallization conditions at pH 6.5, 25 °C and incubation for 10 days before data collection, the Asn109 side chain is intact as supported by the observed electron density.

### QM/MM Metadynamics

#### In silico generation of MjGATase_ASN_109_ for QM/MM studies

SNN formation entails abstraction of a proton from the (n+1) residue backbone amide NH, cyclisation and ammonia loss, a process that appears spontaneous in MjGATase_SNN_109_. The only two structures with an amino acid residue at the n+1 position that yield an intact Asn side chain are those in which the neighbouring residues are mutated. To gain further insights into role of neighbouring residues on succinimide formation, we generated a structure wherein the Asn109 side chain is intact, along with retaining the WT neighbours. Hence, firstly, the SNN residue in the MjGATase_SNN_109_ structure was replaced, using PyMOL, by an Asn residue wherein the backbone dihedral angles continued to remain in the bottom right quadrant, a relatively high energy local conformation in the Ramachandran map. Subsequently, energy minimization and equilibration at constant temperature, pressure, and volume were carried out initially using GROMOS force field [20] followed by one of the standard AMBER family of force fields, AMBER99sb-star-ILDN [35,36]. This structure from classical MD, referred to as MjGATase_ASN_109_ (Fig. 7a), was used for subsequent QM/MM MD metadynamics simulations, with and without the Gaussian bias. Figure S11a-c shows the structural superposition of MjGATase_ASN_109_ on MjGATase_SNN_109_, and the double mutants. Interestingly, the overall structural superposition had similar RMSD values for the mutants and the WT enzyme (0.716 Å, 0.753 Å, and 0.739 Å, respectively for D110V_K151L, K151L_Y158F, and WT), a feature also seen in the loop containing Asn/SNN and in the extended β-sheet region (Fig. S11d-f, h-j). Structural overlay of MjGATase_ASN_109_ on MjGATase_D110V_K151L showed similar disposition of the side chain of Asn109 in the near attack conformation [3,18,32,33] with a van der Waals distance between CG of Asn109 and D110 backbone NH (Fig. S11g). Examination of the classical MD simulation data for the possibility of any of 10 Asn side chains adopting a near attack conformation showed that only Asn109 favoured this conformation with a mean distance of 3 Å (Fig. S9b).

**Figure 7.**
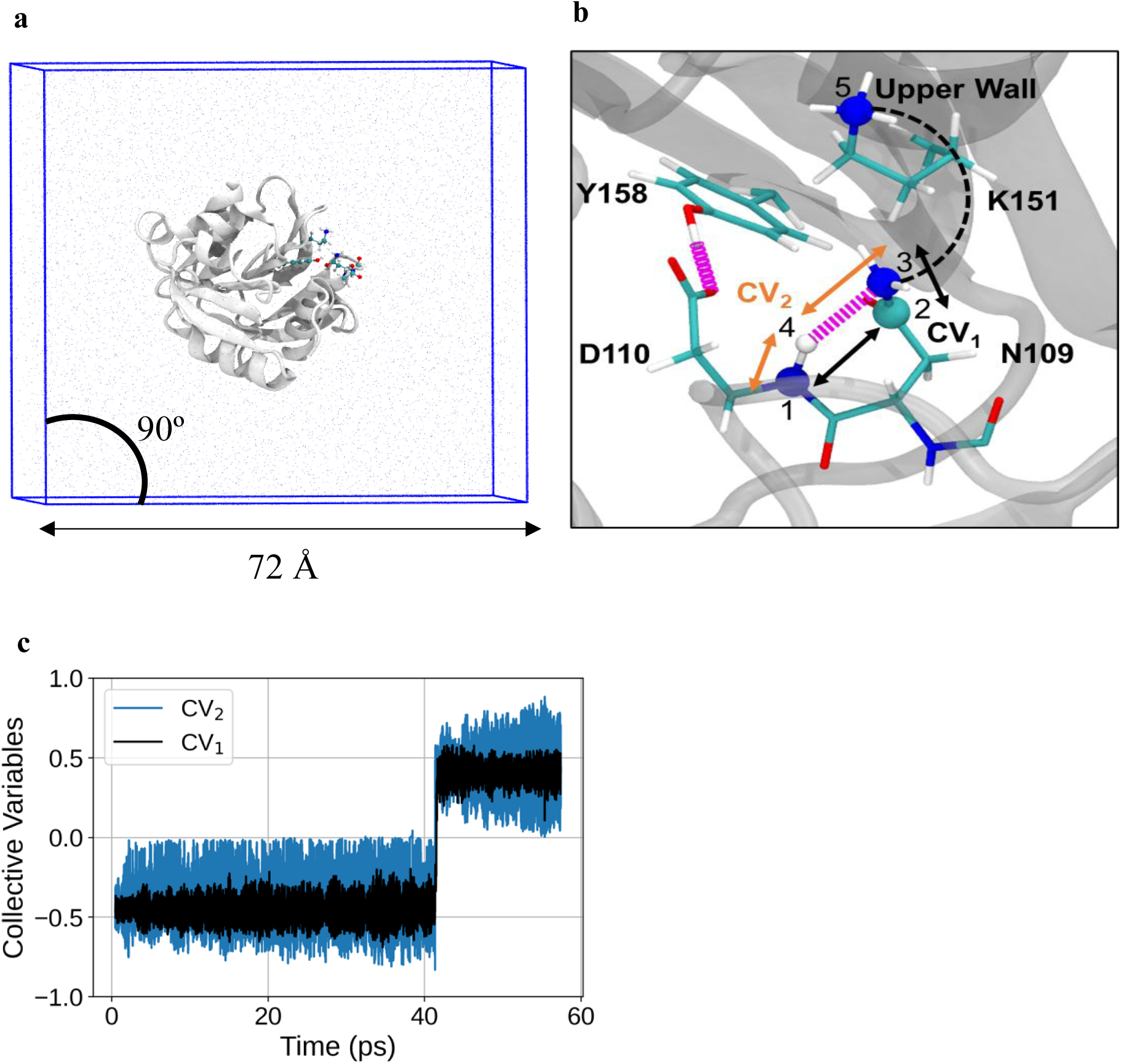
System under computational study, initial conformation of the QM region and time evolution of the collective variables. (a) MjGATase_ASN109 is shown in New Cartoon representation in white. QM region consisting of residues Asn109, Asp110, Lys151, Tyr158, and backbone carbonyl of Glu108 is highlighted in CPK representation. Water molecules are shown as iceblue dots. Ions are not shown for clarity. (b) Zoomed view of the QM region used for QM/MM MD metadynamics simulation. Atoms within QM region are shown in Licorice representation. The remaining protein structure is partly shown in white transparent mode in New Cartoon representation. Water molecules and ions are not shown for clarity. Hydrogen bonds are shown as magenta springs. Atoms involved in the CV definitions and upper wall definition are highlighted in CPK representation. Atoms 1, 2, and 3 are part of CV1 (black arrows) and atoms 1, 3 and 4 are part of CV2 (orange arrows). The upper wall (black dashed line) is between atoms 3 and 5. (c) Time evolution of the two collective variables. CV1 and CV2 show a single transition at around approximately 40.75-41.05 ps.

### Results from QM/MM Metadynamics

To understand the mechanism of SNN formation computationally, we used QM/MM MD metadynamics simulations. The reaction definition included all the three steps – namely, deprotonation, cyclization, and deamidation, represented together through two collective variables. The QM/MM MD run was carried out and monitored for ∼ 57 ps using the initial frame shown in Figure 7b with atoms of Asn109, Asp110, Lys151, Tyr158 and backbone CO of Glu108 in the quantum region and the rest of the protein in the MM region, along with the collective variables CV1 and CV2 as described in methods. Both the CVs are a combination of two coordinates, with CV1 describing the cyclization and deamidation, and CV2 describing the deprotonation and assistance for deamidation. The trajectory yielded only one transition from the reactant basin to the product basin as shown in Figure 7c.

The Ramachandran angles and side chain dihedrals in the structures from the QM/MM run were compared with those of MjGATase_ASN_109_ and MjGATase_SNN_109_ (Fig. S12a-f). Examination of the Ramachandran plots shows that the product states from the simulations are mainly in the bottom left quadrant (Fig. S12c), unlike the experimental product state that is present in the bottom right quadrant (Fig. S12b). However, SNN containing structures extracted from the PDB database display points in both the bottom quadrants of the Ramachandran plot (Fig. S12g). This indicates that due to Asn side chain cyclization, the ψ value is restricted to a range from -105° to -145°, with the dihedral angle φ occurring in both the bottom quadrants (Fig. S12g). In the case of side chain dihedrals of Asn109, Janin plots (Fig. S12d-f, [37]), suggested the partial sampling of the succinimide state (Fig. S12e) in the QM/MM metadynamics trajectory reaction run (Fig. S12f).

To explore the possibility for the (n+1) residue backbone amide hydrogen to bond with any neighbouring basic sites, we calculated the normalized distributions of the corresponding proton abstraction distances. The mutational data presented in the preceding section clearly establish the key role of the proximal carboxylate group of D110 in determining the extent of succinimide formation and earlier studies further implicate D110 in impeding aqueous hydrolysis through a combination of steric and electronic effects [19,20]. The carboxylate group with a pKa of 3.4±1 [38,39] often functions as a base in enzymatic reactions. Figure 8a summarizes the computed distribution of the proton abstraction distance of the D110 carboxylate and also presents comparative plots for the other proximal groups Y158 hydroxyl O, and the Asn109 side chain amide O and N atoms. D110 CG oxygen shows a probability of 0.64 at an abstraction distance of < 2 Å. The proton abstraction distance for D110 CG oxygen was examined as a function of time (Fig. 8b). Interestingly, D110 from early timepoints comes down to a distance of 1 Å but fails to retain the proton. In this context, the possible scenarios are, the proton is transferred back to D110 amide NH or to a water molecule. Interestingly, the highest probability (0.86) at a proton abstraction distance of 1 Å is that of Asn109 amide NG (Fig. 8a), which until 40 ps did not abstract D110 backbone proton (Fig. 8b). However, at around 40 ps, the side chain possibly assumes the near attack conformation by which it directly abstracts the proton and retains it, as seen up to 57 ps (Fig. 8b). D110 CG oxygen deviating away from 1 Å coincides with the Asn109 amide NG having abstracted the proton without exchange between them. Of the two, the more frequent event is D110 carboxylate abstracting the proton.

**Figure 8.**
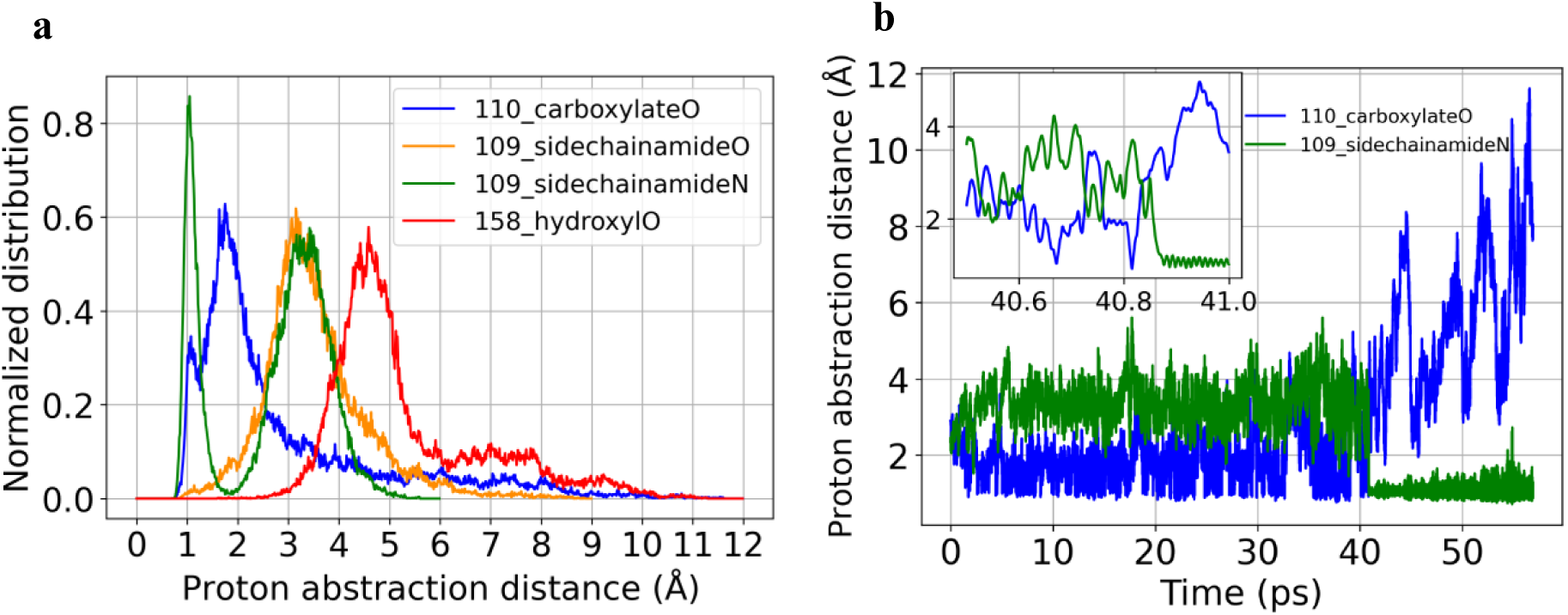
Analysis of the QM/MM MD metadynamics simulation. (a) Normalized distance distributions for the D110 backbone amide proton abstraction by the indicated neighbouring nucleophilic sites. (b) Time evolution of the D110 (n+1) backbone amide proton abstraction by its own side chain carboxylate and by the side chain of Asn109 through the nitrogen atom. Inset shows a zoomed view at time around 40 ps.

The interactions of Asn109 and Asp110 with neighbouring residues were monitored in the classical MD, QM/MM MD, and QM/MM MD metadynamics runs carried out using the energy minimised MjGATase_ASN_109_ structure. The maximum percentage occurrences of contacts in the MD simulation runs that are present in the crystal structures of either MjGATase_SNN_109_ or mutants are described below. The interaction between OD1 of Asn109 and backbone amide hydrogen of Leu111 was found to be retained at a significant level of about 40-45% in the QM/MM MD and QM/MM MD metadynamics runs (Supp. Fig S13a, b). The second interaction between backbone oxygen of Asn109 and backbone amide hydrogen of Phe112 was found to be retained at a much lower extent of 12% and 25% in the QM/MM MD and QM/MM MD metadynamics runs, respectively (Supp. Fig S13c, d). Finally, the contact between the backbone oxygen of D110 residue and backbone amide H of K113 residue was retained in all three simulations to only about 30% (Fig. S13e, f).

Summation of all the deposited hills on the landscape described by the two CVs generated a partial free energy surface, shown in Figure S14a. We extracted a minimum free energy path starting from the reactant state point to an arbitrary point on the landscape. The final point is an arbitrary point, not the product point, because we were unable to fill the product basin significantly, due to limitations in the length of the QM/MM MD metadynamics trajectory. Hence, we proceeded to estimate the activation barrier from the reactant basin. In Figure S14a, the minimum free energy path joining the reactant state (represented by coordinate (-0.4, -0.4) in the CV space) to the arbitrary point (represented by coordinate (0.0, +0.4) in the CV space) is shown as black dashed line and highlights four state points i.e. reactant, 1, 2, and 3. Of the state points 1 to 3, 3 is an arbitrary point *en route* to the product basin (state point 4) and 1 is the barrier between reactant and state point 2. The free energy profile for the segment between states 1 and 2 is shown separately in Figure S14b, where the x-axis corresponds to the number of nodes used in the minimum path-finding algorithm. It provides an activation barrier of ∼3.4 kcal/mol for the deprotonation of the n+1 backbone amide.

To examine these state points, we proceeded to examine the time evolution of the system. We have mainly three parameters to monitor; these are the metadynamics bias and the two CVs, shown in Figure 9. In addition, we had an upper wall bias potential. In Figure 9a, we observe the filling up of the reactant basin by the metadynamics bias around 41 ps. Zooming into the time axis, the inset shows the metadynamics bias along with the two CVs. Figure 9b is a further zoomed view of the same. Here, we see for a stretch of time points, the metadynamics bias is zero and one of the CVs (CV2) crosses zero value in the y-axis while the other (CV1) approaches zero. Time points corresponding to our two state points, 1 and 2 of Figure S14a are shown with vertical dashed green lines in Figure 9b. With progression of time (Fig. 9c), we identified state points 3 and 4 of Figure S14a. At state point 3, CV1 crosses the zero value on the y-axis. To move from state point 2 to 3 and then from 3 to 4, on the free energy surface, the system confronts only small barriers, because we observe that the additional metadynamics bias is fairly low, at a value of ∼0.5 kcal/mol. Near state point 4, both the CV values fluctuate around +0.5, corresponding to the product state.

**Figure 9.**
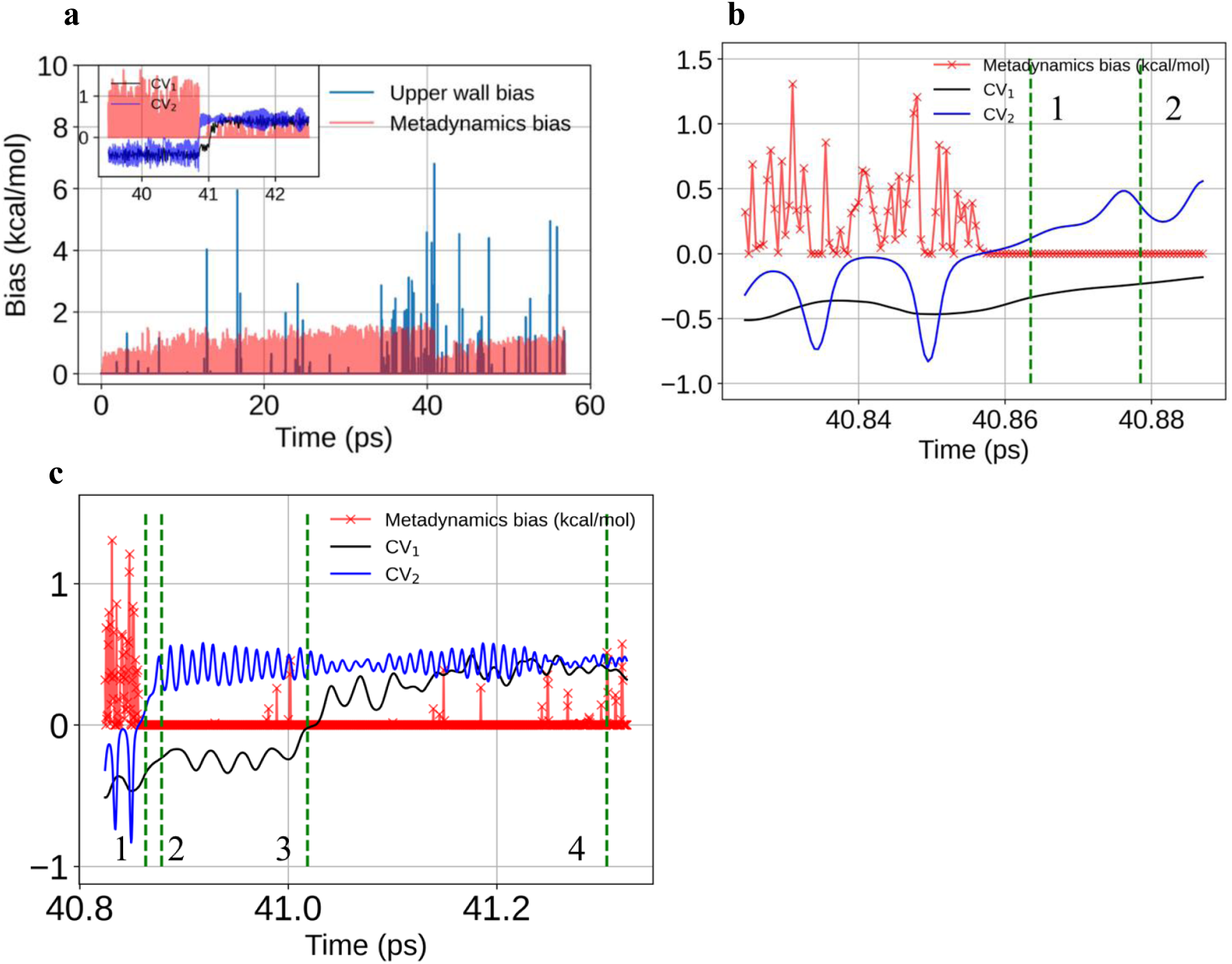
Time evolution of biases and collective variables. (a) Time evolution of the Metadynamics bias and upper wall bias. Inset shows a zoomed view of the time evolution of the two collective variables along with the Metadynamics bias. (b) A further zoomed view of panel (a) near the time point where Metadynamics bias has filled up the reactant basin. State 1 and 2 from Figure S14 (a) are highlighted with dotted vertical green lines. (c) Time advancement towards the product basin. States 1 to 4 from Figure S14 (a) are highlighted with dotted vertical green lines.

Together Figure S14 and Figure 9, provide insights on the free energy surface up to state point 2. Point 1 is the first activated state from the reactant basin with a free energy barrier for the deprotonation step of 3.4 kcal/mol. Since we could not observe product to reactant transition, we do not have a conclusive barrier height for the cyclization and deamidation steps of SNN formation.

Finally, we visualized the structures corresponding to the time points of the four state points (green dashed lines, Fig. 9c) in Figure S14c. State point 1 corresponds to the structure where the backbone amide hydrogen of D110 is detached from the nitrogen and is near the nitrogen nucleus of the side chain amide of Asn109. State point 2 corresponds to the structure where the backbone amide H of D110 now belongs to the side chain amide nucleus of Asn109 completely i.e. C-NH_3_ unit has formed but the loss of ammonia has not happened as the nitrogen is still attached to the side chain amide carbonyl-carbon of Asn109. State point 3 corresponds to the partial cyclization and deamidation, which goes to completion at state point 4, which is the product state.

## Discussion

The spontaneous deamidation of asparaginyl residues in peptides is influenced mainly by the succeeding residue with faster deamidation rates if it is a glycyl residue, and in proteins, the rate is influenced by the primary sequence and three-dimensional structure [11,13,16–18]. Although the t_1/2_ for deamidation of the asparaginyl residue in peptides succeeded by an aspartyl residue is higher when compared to glycine, serine, or alanine at the n+1 position [2,40], in MjGATase, the Asn109 residue succeeded by Asp undergoes spontaneous and complete deamidation leading to the formation of a stable succinimide [19]. This indicates that the three-dimensional architecture of MjGATase plays a major role in the deamidation process supported by the fact that Asn7 succeeded by a Gly residue does not undergo deamidation in the same protein. Our mass spectrometric and crystal structure studies of site-directed MjGATase mutants followed by QM/MM studies provide a mechanistic insight into Asn109 deamidation. The search for mutants having the mass corresponding to unmodified Asn109 gave us the clue that the residues Y158, D110, K113, K151, and E108 have a role in succinimide formation, in decreasing order of their contribution. Further mutational analysis showed that the double mutants, D110V_K151L and K151L_Y158F had the highest proportion of Asn109 intact. Crystal structures of these double mutants confirmed the presence of intact Asn109. Although the single mutant K151L had a negligible population of intact Asn109, the double mutants D110V_K151L and K151L_Y158F had a dramatic increase in the Asn109 intact population. This shows that the 3D environment of the loop harboring SNN/Asn109 is sensitive to local conformational changes with multiple modifications in this environment leading to abrogation of SNN formation. Lysine residues that are spatially proximal to Asn residue can increase the rate of deamidation through stabilization of the negative charge of the transition state before the tetrahedral intermediate formation [41]. This interaction might be facilitated by the field or inductive effect [41]. In MjGATase, K151 and K113 residues that are close in space to Asn109 can play a similar role in favoring SNN formation. Studies on penta/hexapeptides with Asn succeeded by Ser residue established a higher rate of deamidation [2,11,34,40]. In addition, it was observed that Asn or Asp succeeded by Ser residue promoted SNN formation [13,42,43] possibly through the hydroxyl oxygen of the Ser residue H-bonding with the backbone amide or hydroxyl hydrogen making an H-bond with either side chain Asn oxygen or nitrogen atoms [11]. This, in turn, can increase the nucleophilicity of the n+1 residue backbone amide or increase the electrophilicity of the Asn side chain carbonyl carbon, thus aiding in succinimide formation [11,13,42,43].Similarly, in MjGATase, the side chain of Asp110 could be H-bonding with its amide NH, increasing its nucleophilicity. In addition, the hydroxyl group of Y158 that interacts with Asn side chain OD1 and ND2 could lead to an increase in electrophilicity of CG. Additionally, Y158 could be involved in proton relay after D110 abstracts the proton. The residue D/V110 (n+1) in both the double mutant crystal structures and classical MD of MjGATase_ASN_109_ adopted a ψ backbone dihedral value that enhances the backbone amide NH acidity [18,34]. Hence, proton abstraction from the backbone amide succeeding Asn109 is feasible in MjGATase. The n+2 residue can favor the deamidation reaction as observed in triosephosphate isomerase (TPI) [18]. A hydrogen bond interaction between n+2 residue backbone amide and side chain OD1 of Asn71 facilitates the near attack conformation required for SNN formation in TPI [18]. A similar interaction is observed with Asn109 side chain OD1 and L111 (n+2) backbone amide in both the double mutant structures of MjGATase and is also retained after SNN formation. Hence, the n+2 residue could favor SNN formation by positioning Asn in the right orientation.

An optimal conformation of Asn backbone dihedral angle ψ of -120°, side chain dihedrals χ1 of +120°, χ2 of +90° or -90°, and CG_n_-N_n+1_ distance < 3.5 Å enables succinimide formation [3,6,32,33]. Among different conformations seen in MD simulations, adopted by N373 in the P-domain of norovirus capsid protein, a rare *syn* (φ = −180 °and ψ = 0°) backbone conformation of N373 was observed where eclipsing of the N373 nitrogen with the nitrogen of D374 could increase the nucleophilicity of backbone n+1 amide, leading to deamidation [17]. In MjGATase, a *syn* conformation for Asn109 was not observed in the energy-minimized structure of MjGATase_ASN_109_ obtained from classical MD simulations. The N373 residue that undergoes deamidation in Saga P-domain of capsid protein lies on the surface exposed loop and is preceded by a Glu residue and succeeded by an Asp residue [44], is similar to MjGATase sequence E_108_N_109_D_110_ and Asn109 also lies on a loop [19,20]. Despite these similarities, the deamidation is facilitated by different factors in these proteins primarily due to differences in three-dimensional structures.

The QM/MM analysis of the reaction path, using the collective variable description, supports neighbouring group participation, suggesting that proton abstraction from the D110 backbone NH by the side chain carboxylate of D110 is plausible. However, the pathway of this proton beyond its transfer to the carboxylate remains unknown. The only other functional group proximate to D110 NH is the side chain amide of Asn109 as seen in the classical MD simulated structure. The activation barrier for deprotonation is about 3.4 kcal/mol, a low value. From QM/MM and Perturbed Matrix Method (PMM) studies on peptides, with and without including water explicitly, it has been observed that the energy barrier for the overall succinimide formation reaction is in the range of ∼ 20-25 kcal/mol [29,45] or ∼ 30-40 kcal/mol [18,28,33,46,47] respectively, with the former being in agreement with the experimental estimate [6,48]. From earlier work on TPI [18], the energy barrier for the tetrahedral intermediate formation has been shown to be higher at 30-40 kcal/mol. In TPI, unlike MjGATase, SNN formation at Asn71 is a very slow process over days (37.8 days). In contrast, the entire population of recombinant MjGATase isolated from *E. coli* is SNN [19], precluding an easy measurement of the rate of conversion. So, we would expect that the rate for the autocatalysis reaction in MjGATase to be significantly higher than in TPI. Further, as we have not proceeded in our QM/MM analysis beyond deprotonation, it is possible that a subsequent step, either formation of tetrahedral intermediate or ammonia loss could have a higher activation barrier.

In conclusion, we find an agreement in experimental observations on mutants and the QM/MM MD metadynamics simulations. Mass spectrometric analysis of mutants of D110 show that succinimide continues to form, indicating an alternate mode for the removal of the n+1 proton. The near-attack conformation for Asn109, revealed in the QM/MM analysis suggests an alternative path for abstraction of the D110 NH proton, a process that could be enabled by neighbouring Y158 side chain hydroxyl group and the K151 side chain NE amino group.

Our mutational analysis yields an apparently paradoxical result. The K151L single mutant forms the stable succinimide efficiently unlike the Y158F and D110V single mutants that show significantly impaired SNN formation suggesting direct role in autocatalysis and subsequent stabilization of the SNN. Yet in the double mutants the K151L mutation appears to dramatically enhance the impairment of SNN formation in Y158F and D110V single mutants. It is possible that K151 plays a supporting role in SNN formation and stabilization which offsets single mutations at positions Y158 or D110. Interestingly, inspection of a limited dataset of 84 sequences of GATase from archaea reveals an almost complete conservation of K/R151 whereas a very small number of examples containing the Y158F and D/E110G mutations were found [20]. Protein sequences are shaped over evolutionary timescales with selected pressures being driven by biochemical function and structural stability. The cluster of residues D110, Y158, and K151 which are spatially proximal to the stable SNN in MjGATase, present an example of evolutionary selection of a spontaneous post-translational modification that maybe an essential requirement for a protein to remain functional in an organism living in deep-sea hydrothermal vents.

### Methodology

#### Generation of site-directed mutants

The primer pairs with the desired mutation, custom synthesized from Sigma-Aldrich or Eurofins, Bengaluru are listed in Table S3. The MjGATase mutants Y158F, E108Q, K113A, D110V, and D110N were generated by quick change PCR with P1/P2, P19/20, P21/P22, P3/P4 and P23/P24 primers, respectively. The pST39 plasmid with MjGATase gene [19] was used as a template for PCR amplification. DpnI digestion was carried out after PCR and *Escherichia coli* XL10 Gold-competent cells were transformed with the digested DNA. The plasmid was isolated and sequenced to check for the presence of the mutation.

For the generation of E108L, K107L, K151L, E137L, and D110P mutants of MjGATase, overlap PCR was used to generate two amplicons with the primer pairs P15 and RP of the desired mutant, and P16 and FP of the desired mutant using wildtype as a template. The gel-extracted amplicons were used as templates for the generation of the third amplicon using P15/P16 primer pairs, which were also gel-extracted and digested using XbaI and KpnI. pST39 plasmid with MjGATase and MjATPPase genes was used as a vector and restriction digested using XbaI and KpnI. Ligation of the gel-extracted vector and insert with T4 DNA ligase was carried out and *E. coli* DH5α or XL10 Gold competent cells were transformed with the ligated DNA. After screening by PCR, the plasmids from the positive clones were isolated and verified by DNA sequencing for the presence of the mutation.

For the double mutants of MjGATase, overlap PCR was used to generate two amplicons with the primer pairs P4 and P15, and P3 and P16 using MjGATase_K151L as the template for the D110V/K151L mutant and with the primer pairs P18 and P15, and P17 and P16 using MjGATase gene as a template for K151L/Y158F mutant. The gel-extracted amplicons were used as templates for the generation of the third amplicon using P15/P16 primer pairs. pST39 plasmid with MjGATase and MjATPPase genes restriction digested using XbaI and KpnI was used as a vector. The gel-extracted vector and insert were used for AQUA cloning [49] and *E. coli* DH5α competent cells were transformed with the DNA mix. After screening by PCR, the plasmids from the positive clones were isolated and verified by DNA sequencing for the presence of the mutations.

#### Protein expression and purification

*E. coli* Rosetta (DE3) pLysS cells carrying either the pST39_MjGATase or mutant plasmid were grown in 800 ml of Terrific Broth with ampicillin (100 μg ml^-1^) and chloramphenicol (34 μg ml^-1^) till O.D_600_ of 0.4-0.6, induced with 0.3 mM IPTG, and grown for another 3 hours at 37 °C. Thereafter, the culture was centrifuged at 3400 x g for 10 minutes to pellet the cells. Cells resuspended in 20 ml of fresh lysis buffer (20 mM Tris HCl, pH 7.4, 10% glycerol, 0.1 mM EDTA, 2 mM DTT, 0.1 mM PMSF) and lysed using a probe sonicator (Sonics & Materials, Inc). The lysate obtained after sonication was centrifuged at 24000 x g for 45 minutes at 4 °C. The supernatant was collected and heated at 70 °C for 30 minutes to precipitate most of the *E. coli* proteins and centrifuged at 24000 x g for 45 minutes at 4 °C. The supernatant was then treated with 0.01% polyethyleneimine (PEI) to precipitate nucleic acids and centrifuged at 24000 x g for 45 minutes at 4 °C. A Q-Sepharose anion exchange chromatography column connected to a Akta Basic HPLC from GE Healthcare Life Sciences, UK was equilibrated using buffer A (20 mM Tris HCl, pH 8, 10% glycerol, 0.1 mM EDTA, 2 mM DTT, 0.1 mM PMSF), and the supernatant from PEI treatment was loaded on to the column. Thereafter, the column was washed with 3 column volumes of buffer A. The protein was then eluted using a linear gradient of 0 to 30% of 1M NaCl in buffer A. The protein fractions collected were loaded on 12% SDS-PAGE and stained using Coomassie Brilliant Blue. The fractions containing pure protein were pooled, dialyzed, and concentrated using Amicon Ultra 10 kDa cut-off filters. The concentration of the purified protein was estimated by the Bradford method using bovine serum albumin as standard, divided into aliquots, and stored at -80 °C.

#### Mass spectrometric analysis of intact proteins

The MjGATase proteins were diluted to 1 mg ml^-1^ in water and dialyzed against water using a 10 kDa cut-off membrane. The protein was further diluted to 0.1-0.2 mg ml^-1^ in water with 0.1% formic acid. 10-30 pmol of the dialyzed protein sample was injected on to a C8 reverse phase column (Thermo Scientific, 2.1 mm diameter, length: 100 mm, 5 μm particle size, 175Å pore size). Mass spectral analysis was carried out on a Q Exactive HF mass spectrometer (Thermo Scientific, USA) equipped with a Thermo Scientific Dionex Ultimate 3000 UHPLC system at the in-house Central Instrumentation Facility, Molecular Biology and Genetics Unit, JNCASR. The protein was eluted using water containing 0.1% formic acid (solvent A) and acetonitrile containing 0.1% formic acid (solvent B) as the solvent system at a flow rate of 0.3 ml min^-1^. The HPLC run conditions were as follows: equilibration at 10% B for 1 min, gradient from 10% B to 80% B over 6 mins, isocratic at 80% B for 1 min, gradient of 80% B to 95% B in 1 min, isocratic at 95% B for 2 mins, back to 10% B in 1 min and finally equilibrate at 10% B for 3 mins. The column temperature was set to 40 ℃. Full MS scans were acquired at a resolution of 120,000 at m/z 200 Th, scan range of 500 - 2000 m/z, AGC target set to 1e6 with maximum ion injection time of 100 ms. The mass spectrometer was operated at a spray voltage of 3.8 kV, capillary temperature of 320 ℃, auxiliary gas heater temperature of 200 ℃, and S-lens RF level value set at 80. The flow rates of sheath gas and auxiliary gas were set to 20 and 10, respectively.

MjGATase_E108Q protein was diluted to 0.1 mg ml^-1^ in 50% methanol and MS-grade water with 0.1% formic acid. The mass spectrum was acquired by direct injection of the protein through a Hamilton syringe in the positive ion mode. The Q Exactive HF mass spectrometer was operated with a scan range of 400 – 2000 m/z, resolution of 120,000 at m/z 200 Th, spray voltage of 3.5 kV, capillary temperature of 275 °C, probe heater temperature of 200 °C, and S-lens RF level value at 80. The flow rates of sheath gas and auxiliary gas were set to 25 and 10, respectively. The ion injection time and ion target value were set at a threshold of 200 ms and 3e6, respectively.

Data collection and visualization were carried out with Xcalibur v4.1 software and Qual browser (Thermo Scientific). Deconvolution of the spectra was carried out using Biopharma finder^TM^ v3.2 software (Thermo Scientific). Xtract algorithm was used for deconvolution as our data were isotopically resolved. The source spectra were generated by using “Average over selected Retention time” method. A relative abundance threshold cut-off of 10-15% above the noise level of the spectrum was set during deconvolution.

#### In-gel trypsin digestion

The protocol followed was as described earlier [19] and derived from that developed by Mann and co-workers [50]. Briefly, 30 μg of the protein was run on 12% SDS-PAGE and stained with Coomassie Brilliant Blue. The stained gel piece carrying the protein band was cut into small pieces, washed with water, and destained using a 1:1 ratio of methanol and 50 mM Tris HCl, pH 7.0. The gel pieces were dehydrated by first using a 1:1 ratio of acetonitrile and 50 mM Tris HCl, pH 7.0, followed by 100% acetonitrile for 30 seconds. The liquid was removed, and the gel pieces were dried in speed-vac and rehydrated in 25 mM dithiothreitol in 50 mM Tris HCl, pH 7.0, and incubated at 56 ℃ for 20 minutes. Thereafter, the gel pieces were washed with water, dehydrated, and dried as described above. The gel pieces were rehydrated with buffer containing 500 ng of trypsin in 50 mM Tris HCl, pH 7.0 such that it covered the gel pieces and incubated at 37 °C for 12 hours. The tryptic peptides were extracted twice with 50% acetonitrile and 5% formic acid. The extracted peptides were pooled together and dried in a SpeedVac. The dried peptides were dissolved in 2% acetonitrile and 0.1% formic acid and desalted using C18 spin columns (Thermo Scientific). The fractions eluted from the C18 spin columns were dried using a SpeedVac.

#### Mass spectrometric analysis of tryptic peptides

The dried peptides were dissolved in 20 μl of MS-grade water with 0.1% formic acid. Mass spectral analyses were carried out on a Q Exactive HF mass spectrometer (Thermo Scientific, USA) equipped with an Easy nanoLC 1200 (Thermo Scientific). Data were acquired in positive ion mode using the HCD fragmentation method. Easy-Spray column (Thermo Fisher Scientific), PepMap RSLC C18 of 2 μm particle size, 100 Å pore size, 75 μm inner diameter and 25 cm length along with a C18 guard column (Acclaim PepMap100), of 3 µm particle size, 100 Å pore size, 75 µm inner diameter and 2 cm in length was used. Column temperature was set to 40 ℃ and autosampler temperature was set to 4 ℃. Water with 0.1% formic acid as solvent A and 80% acetonitrile with 0.1% formic acid as solvent B was used. The flow rate was set at 300 nl min^-1^.

The peptides of the MjGATase_SNN_109_ and mutants were eluted using the following steps; a gradient of 5% B to 25% B in 65 mins, isocratic at 25% B for 5 mins followed by a gradient of 25% B to 95% B over 15 mins, isocratic at 95% B for 10 mins, gradient of 95% B to 5% B in 2 mins and finally equilibration at 5% B for 8 mins. The mass spectrometer was operated in a data-dependent mode with a scan range of 300 – 3800 m/z and a resolution of 120,000 at m/z 200 Th. The top 15 abundant ions were fragmented by HCD at a normalized collision energy of 30 and the ddMS2 was acquired at a resolution of 15,000 at m/z 200 Th, a scan range of 200 – 2000 m/z, and dynamic exclusion set to 5 sec. The ion injection time and ion target value were set at a threshold of 50 ms and 3e6 for MS scans and 50 ms and 1e5 for MS/MS scans. A nanospray ion source was used with a spray voltage of 1.7 kV, capillary temperature of 275 ℃, and S-lens RF level set to 55. Data acquisition was carried out using Xcalibur software v4.1 (Thermo Scientific).

Data analyses were done with Proteome Discoverer (PD) v2.4 software and Qual browser of Xcalibur software v4.1 (Thermo Scientific). The search engine, Sequest HT in PD was used with an input of MjGATase sequence into the protein database. The following search parameters were used: precursor mass tolerance set to 10 ppm and fragment mass tolerance of 0.05 Da. The maximum missed cleavage sites allowed for tryptic peptides was set to 2. Dynamic modifications of oxidation at Met (+15.995 Da), ammonia-loss at Asn (-17.027 Da) and deamidation at Asn/Gln (+0.984 Da) leading to the conversion of -CONH_2_ to -COOH were included.

#### Crystallisation and data collection

Crystallization of the mutant proteins of MjGATase, D110P, D110_K151L and K151L_Y158F was set up under a 1:1 mixture of silicon and paraffin oil using the micro-batch method. The conditions from commercially available kits of Hampton Research (USA) and Molecular Dimensions (UK) were used to set up the crystallization screens and the trays were kept in a room maintained at 25 ℃. The crystals were obtained within 7-10 days. MjGATase_D110P crystals were obtained in 0.2 M zinc acetate dihydrate with 20 % PEG 3350, pH 6.4., MjGATase_D110V_K151L crystals were obtained in 0.1M MES, pH 6.5, 0.2 M ammonium sulfate, 30 % PEG monomethyl ether 5000, and MjGATase_K151L_Y158F crystals were obtained in 0.1 M MES, pH 6.5, 0.2 M ammonium sulfate, 30 % PEG monomethyl ether 5000. The crystal, prior to diffraction was pre-soaked in the cryoprotectant containing 20 % glycerol for MjGATase_D110P and MjGATase_D110V_K151L, and 10% glycerol for MjGATase_K151L_Y158F before mounting it on the goniometer head.

X-ray diffraction data were collected at the X-ray facility at the Molecular Biophysics Unit, Indian Institute of Science, Bangalore. X-ray diffraction data on all crystals were collected using a Rigaku RU200 X-ray diffractometer (Tokyo, Japan). This was equipped with a rotating anode-type light source with an osmic mirror that gives a monochromatic light source of wavelength 1.54179 Å.

#### Structure determination and refinement

Processing of the diffraction images was carried out using iMOSFLM [51]. SCALA module and PHASER module of CCP4 were used for scaling and phasing, respectively [52–54]. The molecular replacement (MR) method was used to obtain structure solutions using MjGATase_SNN_109_ (PDB ID: 7D40) structure as the model. The refinement of the structures was carried out using the REFMAC module of CCP4 [55] and manual refinement was carried out using COOT software [56]. The structures are available online on the RCSB-PDB website with the PDB IDs as mentioned in Table 1. All structure analyses were carried out using PyMOL [57].

#### Thermal denaturation

The denaturation temperature was determined by absorbance measurements using a UV spectrophotometer. Solutions of MjGATase_SNN_109_ and mutant D110P, D110V_K151L, and K151L_Y158F proteins of MjGATase at 10 μM protein concentration in 20 mM Tris, pH 7.4, were incubated for 30 minutes at different temperatures of 25, 70, 75, 80, 85, and 90 ℃ in a dry bath. At 100 ℃, the proteins were incubated for 15 minutes. The proteins were later centrifuged at 16000 x g for 15 minutes to pellet down the precipitated protein. The supernatant containing the remaining soluble protein was collected and the absorbance spectrum was acquired from 200 – 400 nm on a UV-Vis spectrophotometer (Shimadzu, UV-1780). The spectrum of the buffer that was treated in the same manner as the protein samples was also acquired and subtracted from the protein spectrum. The T_m_ was then estimated from the plot of the absorbance at 280 nm against the temperature.

#### Rate of denaturation

To estimate the rate of denaturation of the MjGATase_SNN_109_ and mutant D110V_K151L and K151L_Y158F proteins of MjGATase, the absorbance spectra were recorded for solutions of the proteins heated at a fixed temperature for varying times. Solutions of WT and mutant proteins at a protein concentration of 10 μM in 20 mM Tris, pH 7.4, were initially equilibrated at 70 ℃ for 7 minutes in a dry bath and ramped up to 80 ℃. At 80 ℃, protein solutions were incubated for 10, 15, 30, 45, 60, 90, 120, and 180 minutes. Thereafter, the solutions were centrifuged at 16000 x g for 15 minutes to pellet down the precipitated protein. The soluble protein in the supernatant was collected and the absorbance spectrum was acquired from 200 – 400 nm on a UV-Vis spectrophotometer (Shimadzu, UV-1780). The protein spectra were corrected for the background by subtracting the spectrum of buffer components. The rate of denaturation was then estimated from the plot of the absorbance at 280 nm against the incubation time.

#### Estimating the increase in SNN population by lowering the pH of protein

MjGATase_D110V_K151L protein at 10 μM concentration in 20 mM ammonium formate, pH 3.0 was incubated at 60 ℃ for 16.5 hours and subjected to LC-MS analysis. The same protein aliquot stored in -80 ℃ was diluted to 10 μM concentration in 20 mM ammonium formate, pH 3.0 and without prior incubation, LC-MS analysis was carried out. This served as the 0^th^ timepoint of the experiment. After incubation, the protein sample was centrifuged at 16000 x g for 10 minutes at 4 ℃ before loading on the column. 5 pmol of the protein sample was injected on to a reverse phase C8 column (Thermo Scientific, 2.1 mm diameter, 100 mm length, 5 μm particle size and 175 Å pore size). Mass spectral analysis was carried out on a Q Exactive HF mass spectrometer (Thermo Scientific, USA) connected to a Thermo Scientific Dionex Ultimate 3000 UHPLC system. The column temperature was maintained at 30 ℃. The injected protein was eluted using water with 0.1 % formic acid (solvent A) and acetonitrile with 0.1% formic acid (solvent B) as the solvent systems and a flow rate of 0.3 ml min^-1^. The gradient used, MS settings, HESI source settings, data collection, visualization, and deconvolution are as mentioned above in mass spectrometric analysis of the intact proteins.

#### Computational methodology

The MjGATase_SNN_109_ contains the PTM, succinimide, at the 109^th^ position (PDB ID: 7D40) [19,20], a product of Asn deamination and cyclization of the side chain CG with the n+1 backbone amide nitrogen. To understand the mechanism of this succinimide formation, we needed to generate the initial configuration or the reactant state (containing Asn) through in-silico mutation. We used chain A from PDB 7D40 and PyMOL [57] for this purpose. The φ and ψ values for succinimide is in the fourth quadrant of the Ramachandran Map, which is a disallowed region for L-amino acids. Hence, Classical molecular dynamics (CMD) was carried out in two steps: first, using a united atom force field GROMOS54A7 [58,59] and second with AMBER99sb-star-ILDNforce field [35,36,60]. By directly using AMBER force field alone, we could not obtain the right set of backbone dihedral angles for Asn109. Upon employing GROMOS force field, backbone dihedral angles for the residue 109 were sampled in the allowed region of the Ramachandran plot. The last frame from the GROMOS trajectory was used as the initial conformation to continue with AMBER family of forcefield. Along with AMBER99sb-star-ILDN for protein, TIP3P [61] parameters were used for water molecules.

We used GROMACS software ([62] http://www.gromacs.org) for classical molecular dynamics (CMD) simulations and CP2K software [63] patched with PLUMED [64,65] for Quantum Mechanics/Molecular Mechanics molecular dynamics (QM/MM MD) simulations. The protein was placed in a periodic cubic box with 20 Å periodic image distance (box length 72 Å) and solvated using the GROMACS solvation module. Protonation states of ionizable residues were set through the GROMACS default program. Three sodium ions were added to neutralize the complete system. Periodic boundary conditions were applied in all three directions. For maintaining temperature and pressure, we used the Bussi-Donadio-Parrinello thermostat [66] and the Parrinello-Rahman barostat [67], respectively. Time constants for temperature coupling at 298 K were 0.5 ps and 100 fs for CMD and QM/MM MD simulations, respectively. Pressure coupling was used only for CMD runs during equilibration. Pressure was maintained at 1 bar isotropically (isothermal compressibility: 4.5x10^−5^ bar^−1^) with a 1.0 ps time constant. Real space cutoffs for the Lennard-Jones and Coulomb interactions were set to 10 Å and 14 Å for AMBER and GROMOS, respectively. Lorentz-Berthelot mixing rules were used to obtain the LJ parameters between two atom types. The particle Mesh Ewald (PME) [68] method with an interpolation order of 4 and a relative tolerance of 10^−5^ was used to calculate the electrostatic interactions for distances above 10 (or 14) Å for CMD calculations in GROMACS. A Smooth particle Mesh Ewald (SPME) [69] method with an interpolation order of 6 and a relative tolerance of 10^−6^ was used to calculate the electrostatic interactions for distances above 10 Å for MM calculations in CP2K. For running the QM/MM MD simulations in CP2K, QUICKSTEP [70] and FIST modules were used. The quantum region was described at the DFT level with the Perdew-Burke-Ernzerhof (PBE) functional [71]. A double-ζ valence-polarized (DZVP) basis set was used along with Goedecker-Teter-Hutter pseudopotentials [72] (DZVPGTH-PBE). The plane wave cutoff was set at 300 Ry, and DFT-D3 dispersion correction was applied [73]. The MM region was described using the same force field used for CMD simulations. The reaction was studied using an enhanced sampling technique (non-tempered metadynamics [74] with two reaction coordinates (i.e. collective variables) and the electrostatic embedding framework to account for the electrostatic interactions between QM and MM regions. Our collective variables (along with the upper wall bias) are shown in Figure 7b in the reactant state generated from the multistep multiscale equilibration. Each of the two collective variables was a combination (using PLUMED function COMBINE) of two coordination numbers (using PLUMED function COORDINATION, see Supplementary text section for the same). CV_1_ is the combination of the number of contacts between “atom 1 and atom 2” and “atom 2 and atom 3”. CV_2_ is the combination of the number of contacts between “atom 3 and atom 4” and “atom 1 and atom 4”. R_0_ value used for the two sets of coordination numbers defining CV_1_, i.e., (1, 2) and (2, 3) was 1.34 Å each, respectively. And the same was 1.00 Å each for defining CV_2_, i.e., (3, 4) and (1, 4). Default values were used for other parameters. So, as per the definition, the values for the combined coordination numbers or collective variables of the reactant and product states were (-0.5, -0.5) and (+0.5, +0.5), respectively (Fig. 7a). In addition, we used an upper wall bias (using PLUMED function UPPER WALL) between the two nitrogen atoms (side chain amide-N of Asn109 and side chain amino-N of Lys151, labelled as atom 3 and atom 5, respectively) at 5.5 Å with a force constant of 200 kcal/mol (remaining parameters with default values). For the metadynamics bias, the sigma values for CV_1_ and CV_2_ were 0.005 and 0.006, respectively and 0.59 kcal/mol hills were added every 100 fs. During analysis, the criteria for successful hydrogen bonding were fixed at a donor-acceptor distance of < 3.5 Å and a donor-H-acceptor angle of >140°. The bin width for histogramming is chosen as .01 Å.

Time steps for integrating the equations of motion for CMD and QM/MM MD simulations were set as 1 and 0.5 fs, respectively. Leap-frog algorithm was used for solving equations of motion. Position restraining was applied using a harmonic potential with a force constant of 10^3^ kJmol^−1^nm^−2^ for the non-hydrogen atoms. All bonds were constrained using LINCS with order 4 and a warn angle of 30^◦^.

The general protocol for equilibration using both forcefields for CMD consisted of four steps: a) energy minimization of water molecules (position restraining the protein), b) energy minimization of protein atoms (position restraining water), c) equilibration at constant volume and temperature (NVT ensemble) by raising the temperature from 0 to 298 K over 500 ps and keeping at 298 K for the next 500 ps, and d) equilibration at constant temperature and pressure (NPT ensemble) at 298 K and 1 atm, respectively for 1 ns. This was followed by the production run in NVT ensemble at 298 K for 100 ns. The non-hydrogen protein coordinates from the 100^th^ ns frame of the GROMOS trajectory was used as the initial conformation to continue with AMBER force field. From the AMBER production trajectory, one arbitrary protein conformation was used to initiate QM/MM MD simulations (Fig. 7b).

We have carried out one CMD simulation with MjGATase_SNN_109_ structure as well. This consisted of three steps of equilibration – i) energy minimization, ii) equilibration at constant volume and temperature (NVT ensemble), and iii) equilibration at constant temperature and pressure (NPT ensemble) which is followed by a production run in NVT. The same GROMOS parameter set was used here as well.

Treating the complete system with QM description is highly computationally expensive for such a large system; hence, only some residues relevant to the reaction (suggested from experiments) were considered at the QM level and others were treated at the MM level. Hence, the QM level contained residues 109 (Asn), 110 (Asp), 151 (Lys), and 158 (Tyr). To make sure that we were not cutting through any polarizable bonds, a part of residue 108 (backbone carbonyl- of Glu) were included in the QM region. The total number of quantum atoms was thus 61 (Fig. 7b) that included four link atoms (Hydrogen atoms) capping the QM region. The quantum box spanned 15 Å in three directions. QM/MM equilibration was done in three steps- a) energy minimization with mechanical embedding, b) energy minimization with electrostatic embedding, and c) equilibration at constant volume and temperature (NVT ensemble) with electrostatic embedding.

A summary of simulations is tabulated in Table S4. The complete system is shown in Figure 7a.

Note: Simulation input files are available at https://github.com/Oishika-1/QM-MM_MjGATase_Succinimide. Simulation data files and analysis scripts would be shared upon request. The reaction VMD movie has been provided at https://drive.google.com/drive/u/0/folders/1dmEbYcDjfDQnFDsQDvq_IthEq4mt9vIa.

## CRediT authorship contribution statement

**Anusha Chandrashekarmath:** Data curation, Formal analysis, Investigation, Methodology, Validation, Visualization, Writing – original draft, Writing – review and editing. **Oishika Jash:** Data curation, Formal analysis, Investigation, Methodology, Validation, Visualization, Writing – original draft, Writing – review and editing. **Karandeep Singh:** Investigation. **Asutosh Bellur:** Investigation. **Chitralekha Sen Roy:** Investigation. **Aparna Dongre:** Investigation. **Nayana Edavan Chathoth:** Methodology. **Padmesh Anjukandi:** Methodology. **Sanjeev Kumar:** Investigation. **Souradip Mukherjee:** Investigation. **Padmanabhan Balaram:** Formal analysis, Supervision, Writing – original draft, Writing – review and editing. **Sundaram Balasubramanian:** Conceptualization, Data curation, Formal analysis, Funding acquisition, Methodology, Project administration, Resources, Supervision, Validation, Visualization, Writing – original draft, Writing – review and editing. **Hemalatha Balaram:** Conceptualization, Data curation, Formal analysis, Funding acquisition, Methodology, Project administration, Resources, Supervision, Validation, Visualization, Writing – original draft, Writing – review and editing.

## Acknowledgements

We are grateful for the support and resources provided by ‘PARAM Yukti’ facility under the National Supercomputing Mission, Government of India, at the Jawaharlal Nehru Centre for Advanced Scientific Research (JNCASR). O.J. and A.C. thank Council of Scientific and Industrial Research and Department of Biotechnology, India, respectively for the research fellowship. N.C is funded by ERG Project funds (2024-238-CHY-PAA-ERG-SP) of Dr. Padmesh A from IIT Palakkad. This work was also supported by grants from the Department of Biotechnology, India (Project no. BT/INF/22/SP27679/2018). HB thanks JC Bose fellowship for funding (SR/S2/RNJ-JCB2009). Intramural funding from JNCASR is acknowledged. O.J. thanks Anand Srivastava for discussions on metadynamics, Sudip Das for help with QM/MM MD simulation protocol and Sudarshan Behera for discussions at the beginning of the project. We thank the Molecular Biophysics Unit, Indian Institute of Science for the use of the X-ray diffractometer. The use of Mass spectrometry facility at Jawaharlal Nehru Centre for Advanced Scientific Research is acknowledged.

