## Supplement for "Insights into the mechanism of succinimide formation in an archaeal glutaminase"

### Supplementary text

For defining the collective variables, we have used COORDINATION and COMBINE functions from PLUMED (https://www.plumed.org/doc-v2.9/user-doc/html/_c_o_o_r_d_i_n_a_t_i_o_n.html and https://www.plumed.org/doc-v2.9/user-doc/html/_c_o_m_b_i_n_e.html, respectively). Each collective variable in a linear combination of two coordination numbers. Parameter values should be referred from Github input files. Following is the functions used:

The coordination number definition:

$$cn1 = cn2 = cn3 = cn4 = \frac{\left(1 - \frac{rij}{ro}\right)^6}{\left(1 - \frac{rij}{ro}\right)^{12}}$$

And the combination definition:

$$c1 = cn1 - cn2$$

$$c2 = cn3 - cn4$$

For upper wall bias (https://www.plumed.org/doc-v2.9/user-doc/html/_u_p_p_e_r__w_a_l_l_s.html), the definition is:

$$uw = k\,(x - a)^2$$

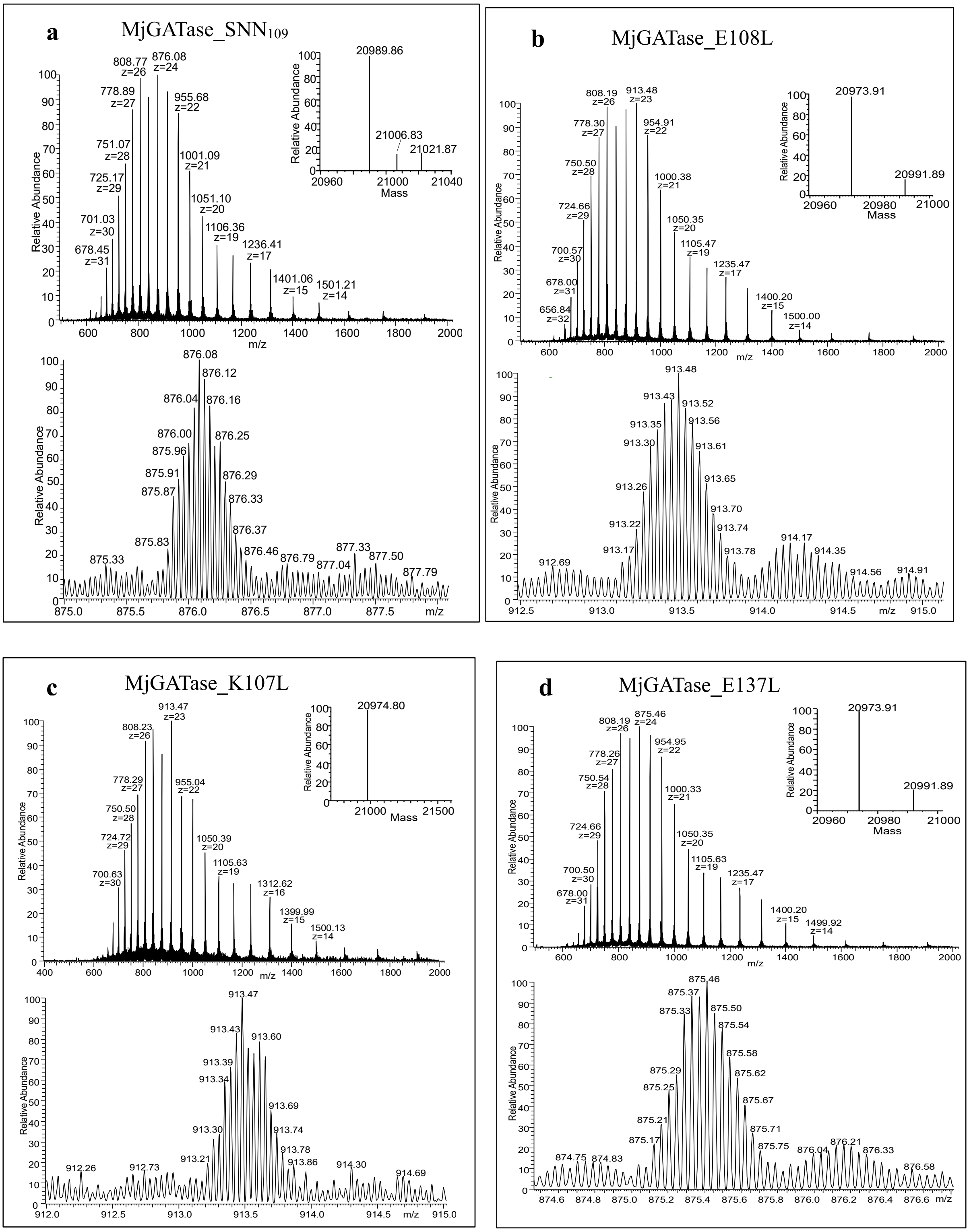
a
MjGATase_SNN109
20989.86
21006.83
21021.87
Mass
Relative Abundance
m/z
b
MjGATase_E108L
20973.91
20991.89
c
MjGATase_K107L
20974.80
d
MjGATase_E137L
20973.91
20991.89

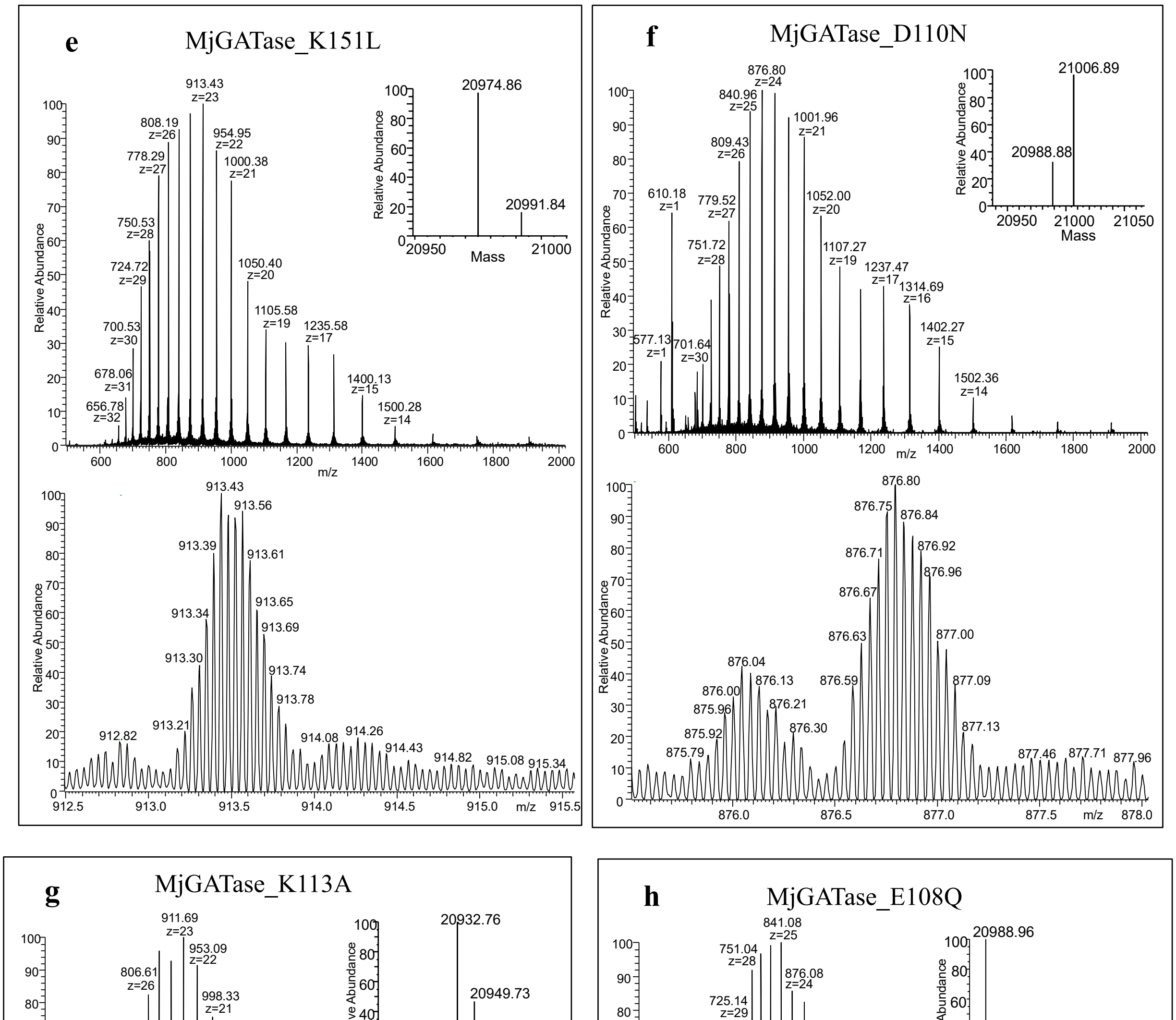


**Supplementary Figure S1. Mass spectra of WT and mutants of MjGATase**. In each main panel, the top sub panel shows the entire mass spectrum of the protein with the inset showing the monoisotopic mass obtained after deconvolution. The bottom panel shows the expanded spectrum of a single charge state. (a) LC-MS of

MjGATase_$SNN_{109}$ ($M_{calc}$ 21006.88 Da). Deconvoluted spectrum in the inset shows a major population with $M_{obs}$ of 20989.86 Da corresponding to SNN109 and, two small populations of 21006.83 Da and 21021.87 Da corresponding to the expected mass of the protein with Asn109 and a methionine oxidised species containing SNN109, respectively. (b) LC-MS of MjGATase_E108L ($M_{calc}$ 20990.92 Da). Deconvoluted spectrum in the inset shows a major population with $M_{obs}$ 20973.91 Da corresponding to SNN109 and a small population having 20991.89 Da corresponding to SNN109 hydrolysed to D/isoD109. (c) LC-MS of MjGATase_K107L ($M_{calc}$ 20991.87 Da). Deconvoluted spectrum in the inset shows a major population with $M_{obs}$ 20974.8 Da corresponding to SNN109. (d) LC-MS of MjGATase_E137L ($M_{calc}$ 20990.92 Da). Deconvoluted spectrum in the inset shows a major population with $M_{obs}$ 20973.91 Da corresponding to SNN109 and a small population having 20991.89 Da corresponding to SNN109 hydrolysed to D/isoD109. (e) LC-ESI-MS of MjGATase_K151L ($M_{calc}$ 20991.87 Da). Deconvoluted spectrum in the inset shows a major population with $M_{obs}$ of 20974.86 Da corresponding to SNN109 and a small population having a of mass 20991.84 Da corresponding to the expected mass with Asn109 intact. (f) LC-MS of MjGATase_D110N ($M_{calc}$ 21005.9 Da). Deconvoluted spectrum in the inset shows a major population with $M_{obs}$ of 21006.89 Da corresponding to hydrolysed SNN109 leading to D/isoD109 and a small population with mass of 20988.88 Da corresponding to SNN109. (g) LC-MS of MjGATase_K113A ($M_{calc}$ 20949.82 Da). Deconvoluted spectrum in the inset shows a major population with $M_{obs}$ of 20932.76 Da corresponding to SNN109 and a small population having a mass of 20949.73 Da corresponding to the expected mass with Asn109. (h) MS of MjGATase_E108Q ($M_{calc}$ 21005.89 Da). Deconvoluted spectrum in the inset shows a major population with $M_{obs}$ of 20988.96 Da corresponding to SNN109 and minor populations having masses of 21005.91 Da, 21020.95 Da, and 21086.92 Da corresponding to the expected mass with Asn109, methionine oxidised species containing SNN109, and an unassigned molecular species which is 81 Da higher than the expected mass, respectively.

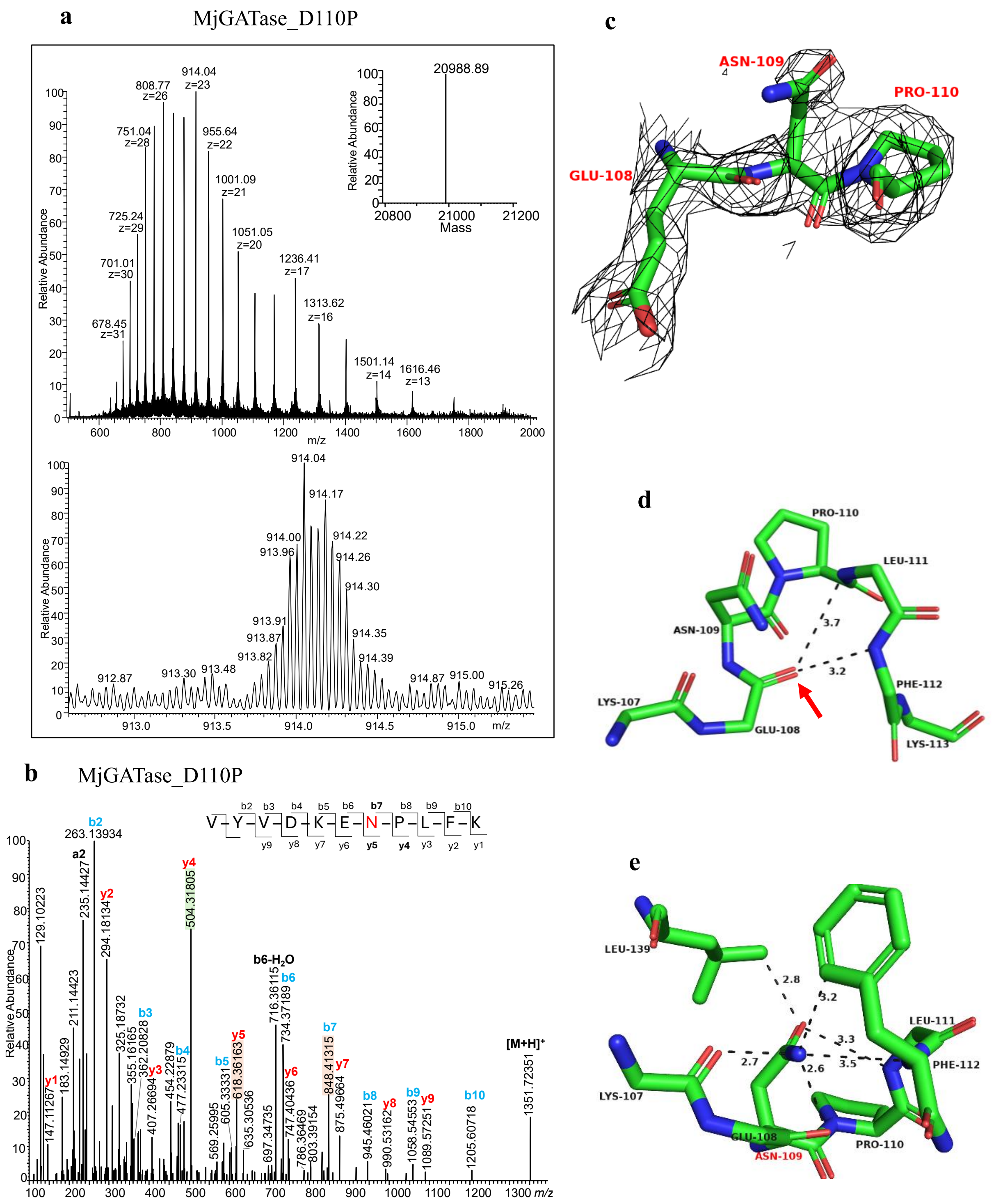


**Supplementary Figure S2. Mass spectrometric and crystal structure analysis of MjGATase_D110P.** (a) LC-MS of MjGATase_D110P ($M_{calc}$ 20988.91 Da). The top panel shows the entire mass spectrum of the protein with the inset showing the monoisotopic mass obtained after deconvolution. The bottom panel shows the expanded the spectrum of a single charge state. The deconvoluted spectrum in the inset shows a mass of 20988.89 Da that is same as the expected mass. (b) MS/MS of peptide of m/z 1351.723 obtained from in-gel tryptic digest of MjGATase_D110P shows the presence of intact Asn109. The y5 and b7 ions with m/z values of 618.36 and 848.41, respectively are highlighted and confirm Asn109. The highlighted y4 ion of m/z 504.31 also confirms the mutation of Asp to Pro at residue 110. (c) The $2F_o$-$F_c$ electron density map contoured to 0.8σ (black mesh) for residues 108, 109, and 110 in the structure of MjGATase_D110P. (d) Structure of Asn109-containing-loop from residues 107-113. The α-turn and β-turn as observed in WT are retained in MjGATase_D110P. The backbone CO of E108 points inwards into the loop similar to WT as indicated by an arrow. (e) Contacts of Asn109 at 4 Å distance cut-off in the structure of MjGATase_D110P.

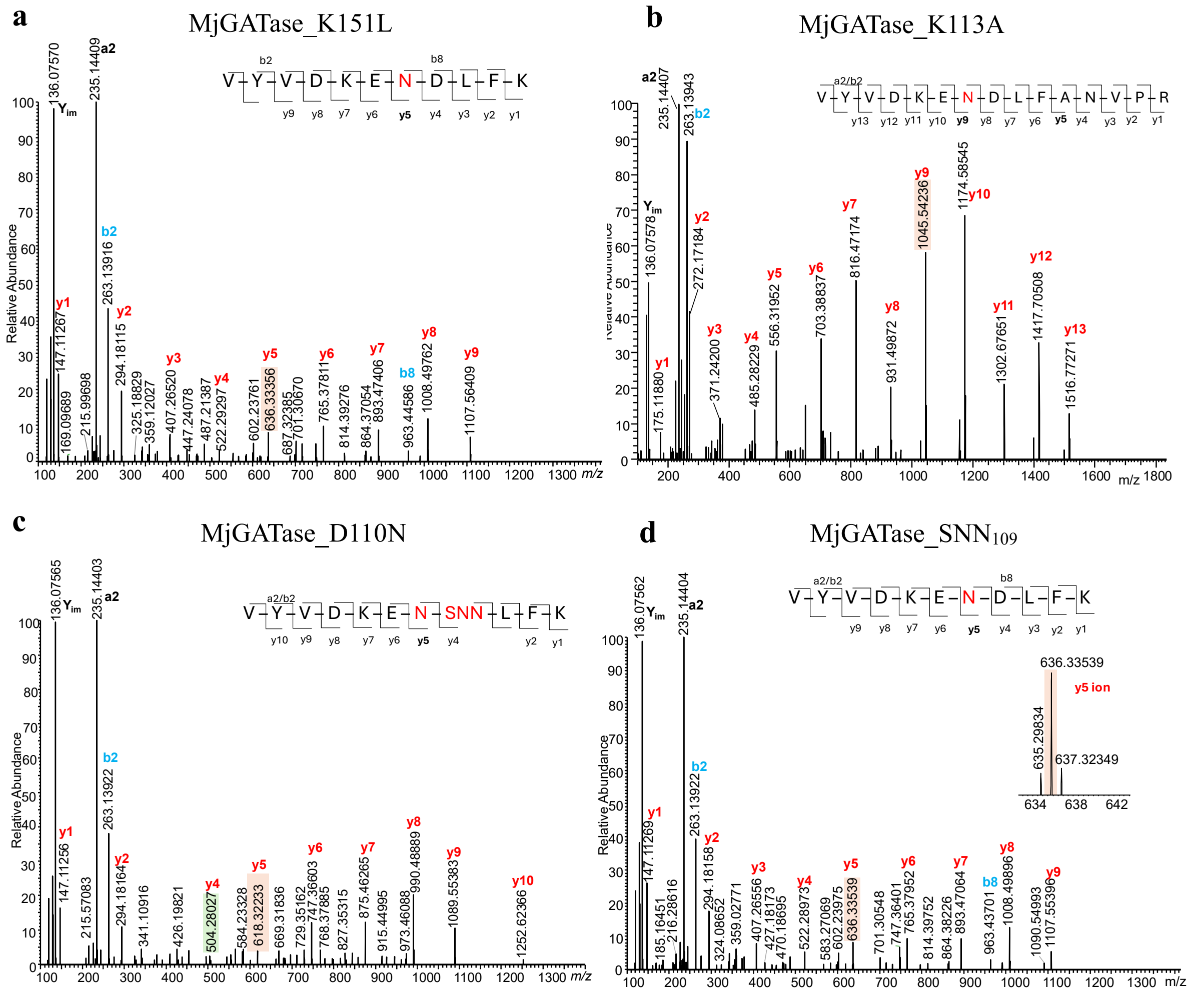


**Supplementary Figure S3**. **MS/MS spectra of the peptides generated by in-gel trypsin digestion of WT and mutants of MjGATase.** (a) MS/MS of the doubly charged peptide of m/z 685.35 obtained from in-gel trypsin digest of MjGATase_K151L shows the presence of intact Asn109. The y5 ion of Asn109 is highlighted. (b) MS/MS of doubly charged peptide of m/z 889.94 obtained from in-gel trypsin digest of MjGATase_K113A shows the presence of intact Asn109. The y9 ion of m/z 1045.54 that confirms Asn109 is highlighted. The y5 ion of m/z 556.31 also confirms the mutation of Lys to Ala at residue 113. (c) MS/MS of the doubly charged peptide of m/z 676.34 obtained from in-gel tryptic digest of MjGATase_D110N shows the presence of intact Asn109. The y5 ion of Asn109 and y4 ion of SNN110 are highlighted. It should be noted that in this mutant the percentage of SNN at the 109 residue is 81% (of a sum of D109, SNN109, and D109SNN110) while deamidation of N110 is a total of 20% (of a sum of D110, SNN110, and D109SNN110) (d) MS/MS of the doubly charged peptide of m/z 685.35 obtained from in-gel tryptic digest of MjGATase_$SNN_{109}$ shows the presence of intact Asn109. The y5 ion of Asn109 is highlighted and the inset shows the isotope distribution of the y5 ion. MS/MS spectra of corresponding peptides with SNN109 or D/isoD109 are provided in Fig. S4e and f for MjGATase_K151L and Fig. S5 for MjGATase_K113A, MjGATase_D110N, and MjGATase_$SNN_{109}$.

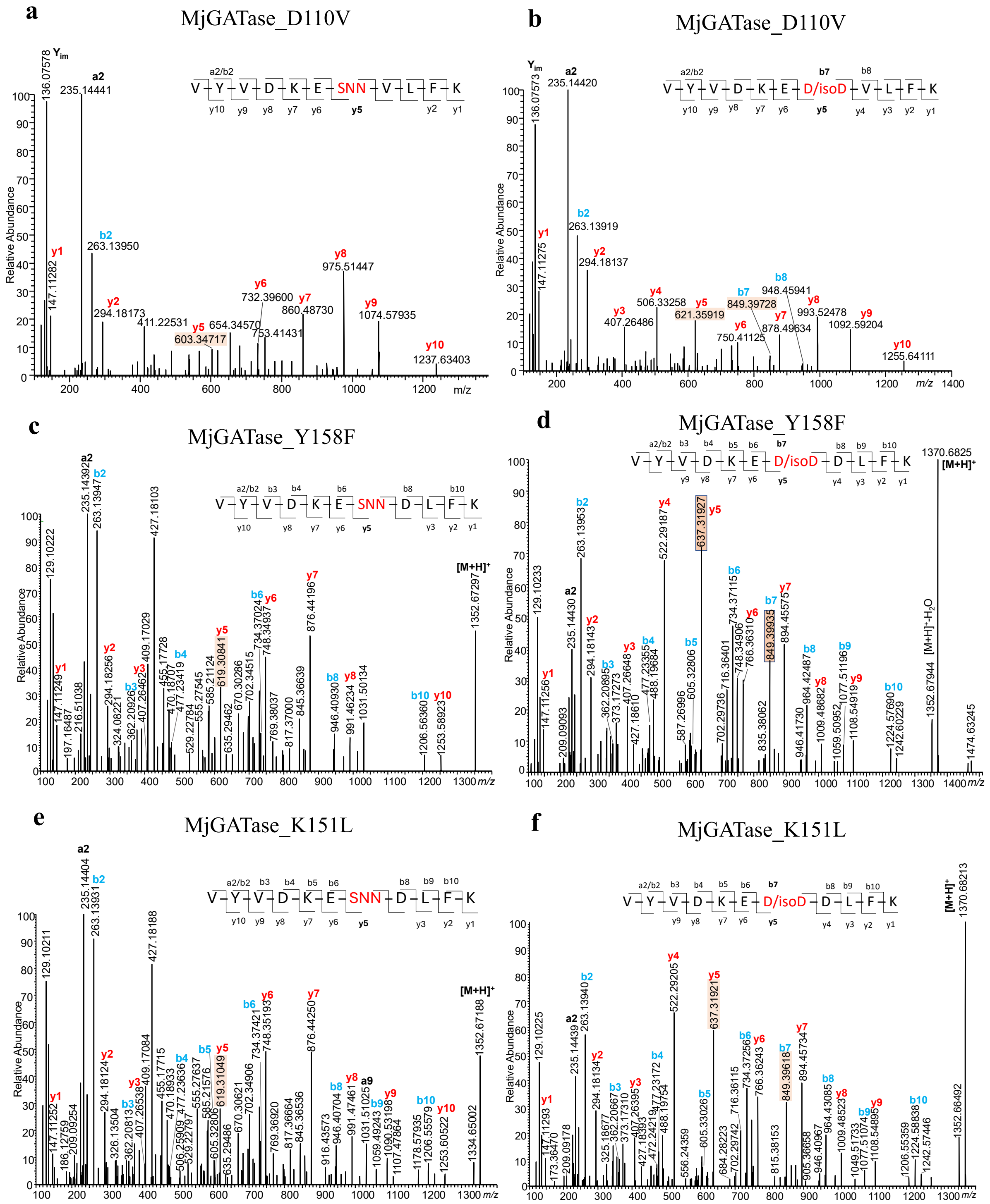


**Supplementary Figure S4. MS/MS spectra of peptides generated by in-gel trypsin digestion of mutants of MjGATase.** The fragment ions are indicated. (a) MS/MS of the doubly charged peptide of m/z 668.86 obtained from in-gel trypsin digestion of MjGATase_D110V. The y5 ion of SNN109 is highlighted. (b) MS/MS of the doubly charged peptide of m/z 677.86 obtained from in-gel trypsin digestion of MjGATase_D110V. The y5 and b7 ions of D/isoD109 are highlighted. (c) MS/MS of the peptide of m/z 1352.67 obtained from in-gel trypsin digestion of MjGATase_Y158F. The y5 ion of SNN109 is highlighted. (d) MS/MS of peptide of m/z 1370.68 obtained from in-gel trypsin digestion of MjGATase_Y158F. The y5 ion and b7 ions of D/isoD109 are highlighted. (e) MS/MS of peptide of m/z 1352.67 obtained from in-gel tryptic digestion of MjGATase_K151L. The y5 ion of SNN109 is highlighted. (f) MS/MS of the peptide of m/z 1370.68 obtained from in-gel trypsin digestion of MjGATase_K151L. The y5 ion and b7 ions of D/isoD109 are highlighted.

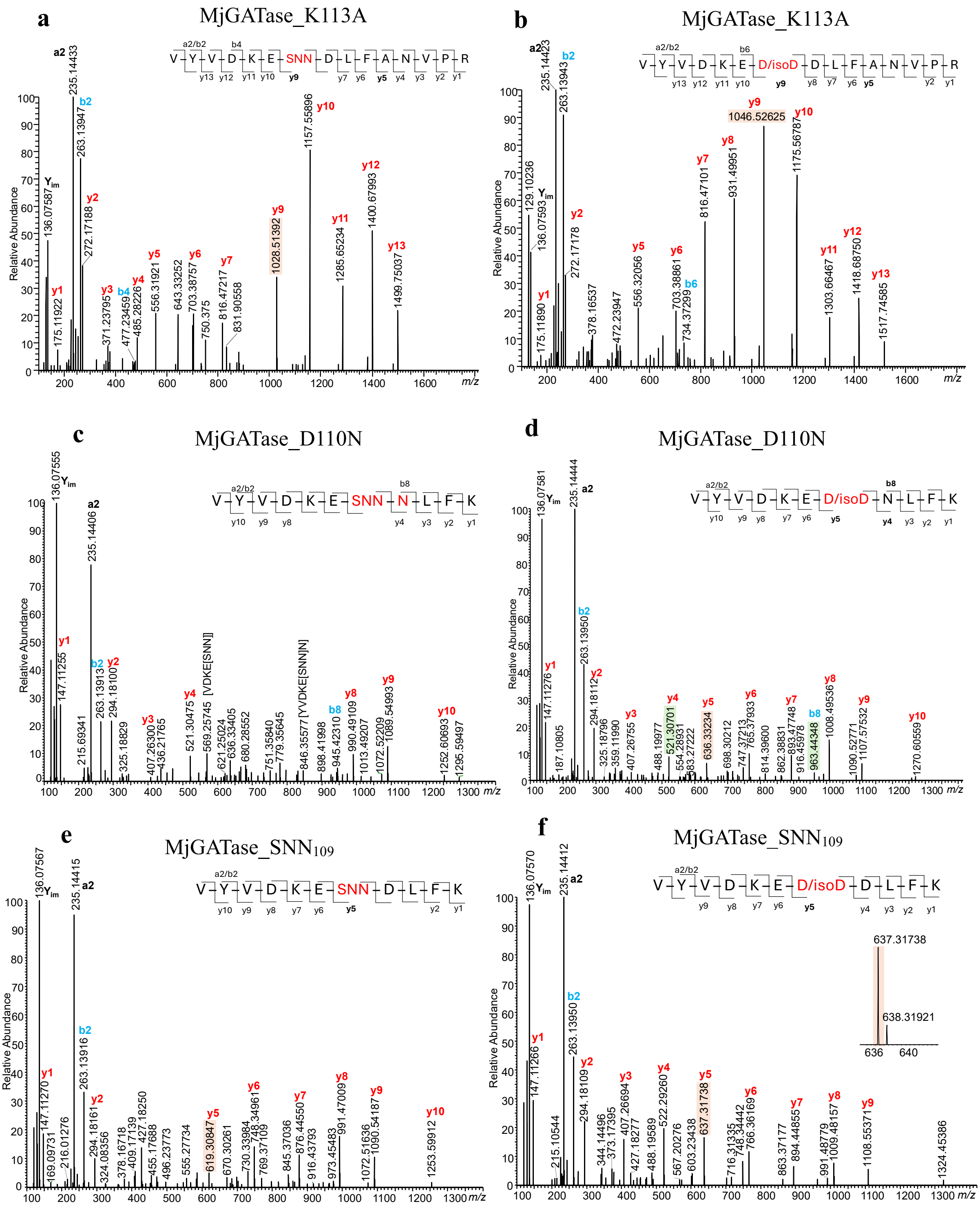


**Supplementary Figure S5. MS/MS spectra of in-gel trypsin digested peptides of MjGATase_SNN$_{109}$ and mutants of MjGATase.** The fragment ions are indicated. (a) MS/MS spectrum of the doubly charged peptide of m/z 881.45 obtained from in-gel trypsin digestion of MjGATase_K113A. The y9 ion of SNN109 is highlighted. (b) MS/MS spectrum of the doubly charged peptide of m/z 890.44 obtained from in-gel trypsin digestion of MjGATase_K113A. The y9 ion of D/isoD109 is highlighted. The y5 ion in (a) and (b) confirm the mutation of Lys to Ala at residue 113. (c) MS/MS spectrum of the doubly charged peptide of m/z 676.34 obtained from in-gel trypsin digestion of MjGATase_D110N. (d) MS/MS spectrum of the doubly charged peptide of m/z 685.35 obtained from in-gel trypsin digestion of MjGATase_D110N. The y5 ion of D/isoD109 is highlighted. The highlighted y4 and b8 ions in (d) and y4 and b8 ions in (c) confirm the mutation of Asp to Asn at residue 110. (e) MS/MS spectrum of the doubly charged peptide of m/z 676.84 obtained from in-gel trypsin digestion of

MjGATase_SNN109. The y5 ion of SNN109 is highlighted. (f) MS/MS spectrum of the doubly charged peptide of m/z 685.84 obtained from in-gel trypsin digestion of MjGATase_SNN109. The y5 ion of D/isoD109 is highlighted and the inset shows the isotope distribution of y5 ion.

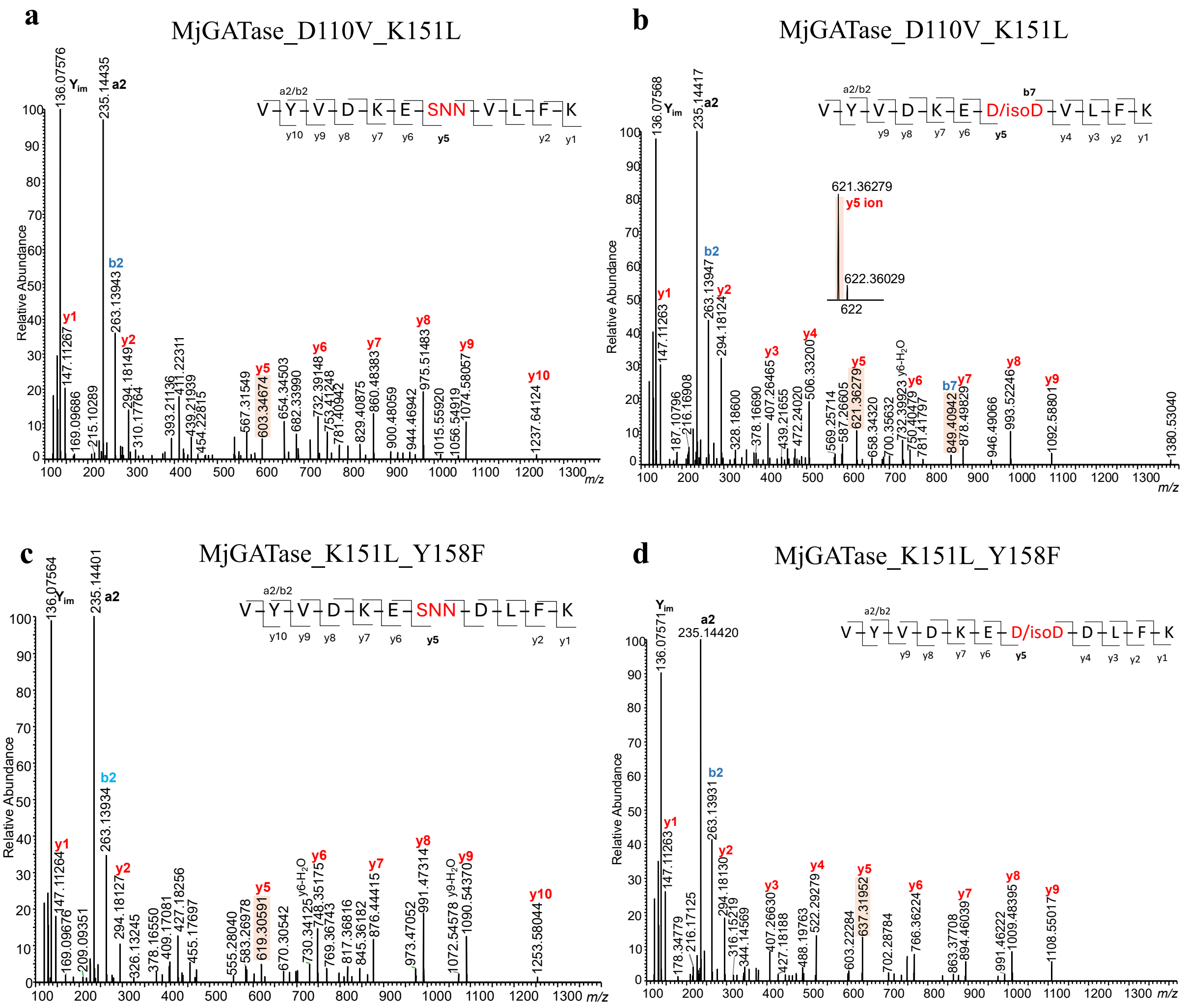


**Supplementary Figure S6. MS/MS spectra of the peptides generated by in-gel trypsin digestion of the double mutants of MjGATase.** The fragment ions are indicated. (a) MS/MS of doubly charged peptide of m/z 668.86 obtained from in-gel trypsin digest of MjGATase_D110V_K151L. The y5 ion of SNN109 is highlighted. (b) MS/MS of doubly charged peptide of m/z 677.85 obtained from in-gel trypsin digest of MjGATase_D110V_K151L. The y5 and b7 ions of D/isoD109 are highlighted and the inset shows the isotope distribution of y5 ion. (c) MS/MS of doubly charged peptide of m/z 676.83 obtained from in-gel trypsin digest of MjGATase_K151L_Y158F. The y5 ion of SNN109 is highlighted. (d) MS/MS of doubly charged peptide of m/z 685.85 obtained from in-gel trypsin digest of MjGATase_K151L_Y158F. The y5 ion of D/isoD109 is highlighted.

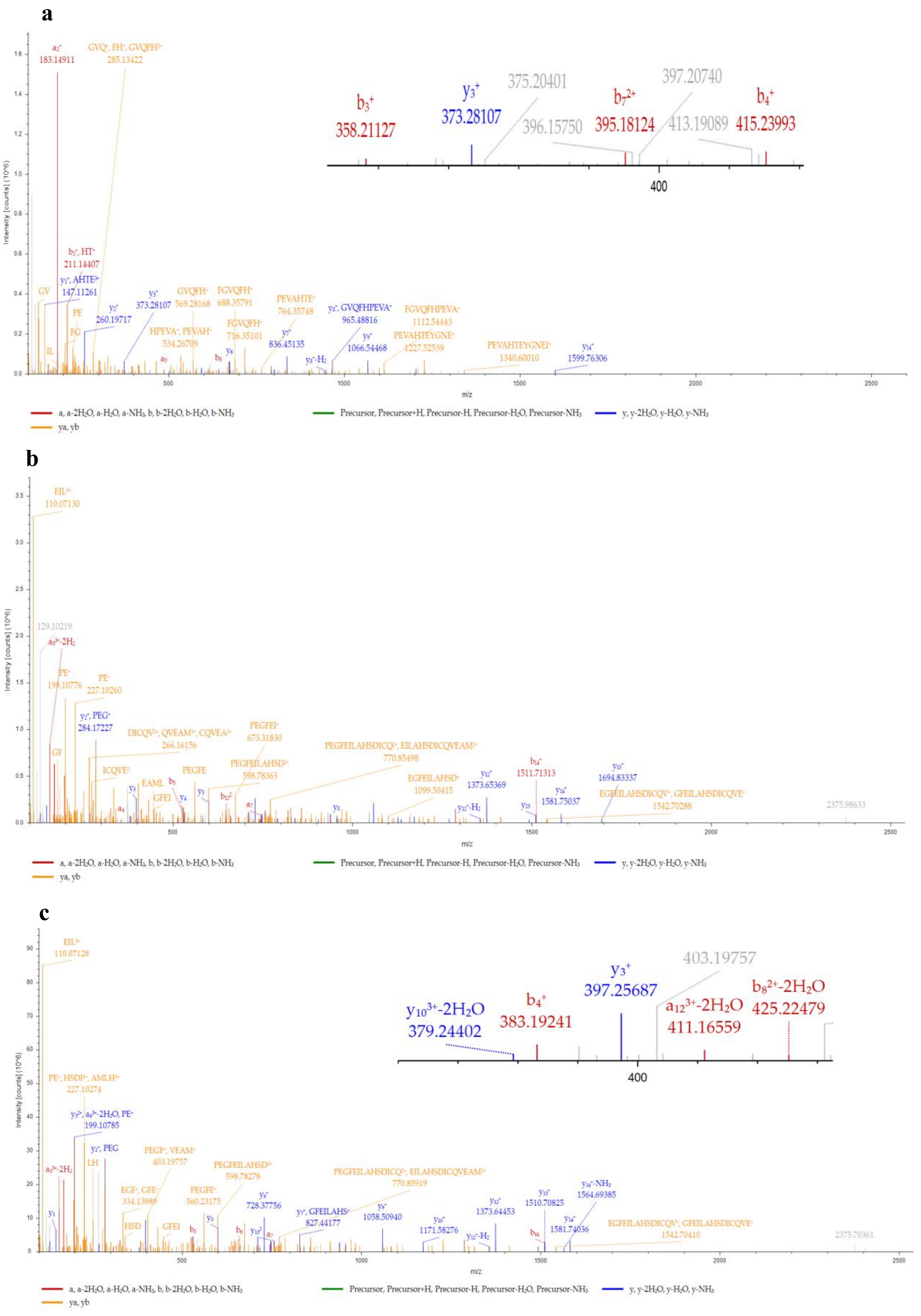


**Supplementary Figure S7. MS/MS analysis of in-gel trypsin digested peptides confirms the mutations in the double mutants of MjGATase.** (a) MS/MS spectrum of the peptide of sequence $PIF_{158}GVQFHPEVAHTEYGNEILK$ of m/z 632.07 (+4) obtained from in-gel trypsin digestion of

MjGATase_K151L_Y158F. The inset shows b3 ion of m/z 358.21127 confirming the mutation of Tyr158 to Phe. (b) MS/MS spectrum of the peptide of sequence VPEGFEILAHSDICQVEAML$_{151}$HK of m/z 822.74 (+3) obtained from in-gel trypsin digestion of MjGATase_K151L_Y158F. The presence of y3 ion confirms the mutation of Lys151 to Leu. (c) MS/MS spectrum of the peptide of sequence VPEGFEILAHSDICQVEAML$_{151}$HK of m/z 822.74 (+3) obtained from in-gel trypsin digestion of MjGATase_D110V_K151L. The inset shows y3 ion of m/z 397.25687 confirming the mutation of Lys151 to Leu. The inset in (a) and (c) is zoomed in x-axis view to show the b/y ions of interest.

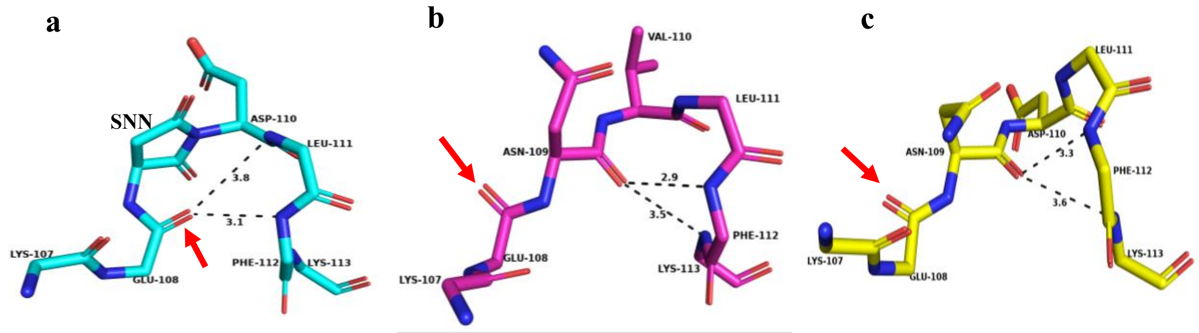


**Supplementary Figure S8. Structural comparison of the SNN/Asn109 loop in WT and the two double mutants**. The structure of the SNN109/Asn109 containing loop from residues 107-113 in (a) MjGATase_SNN$_{109}$, (b) MjGATase_D110V_K151L and (c) MjGATase_K151L_Y158F. An α-turn and a β-turn can be seen in the MjGATase_SNN$_{109}$ structure (a). Abrogation of the SNN induced α-turn is observed in MjGATase_D110V_K151L (b) and MjGATase_K151L_Y158F (c) structures. The orientation of backbone E108 CO indicated with an arrow is different in double mutant structures.

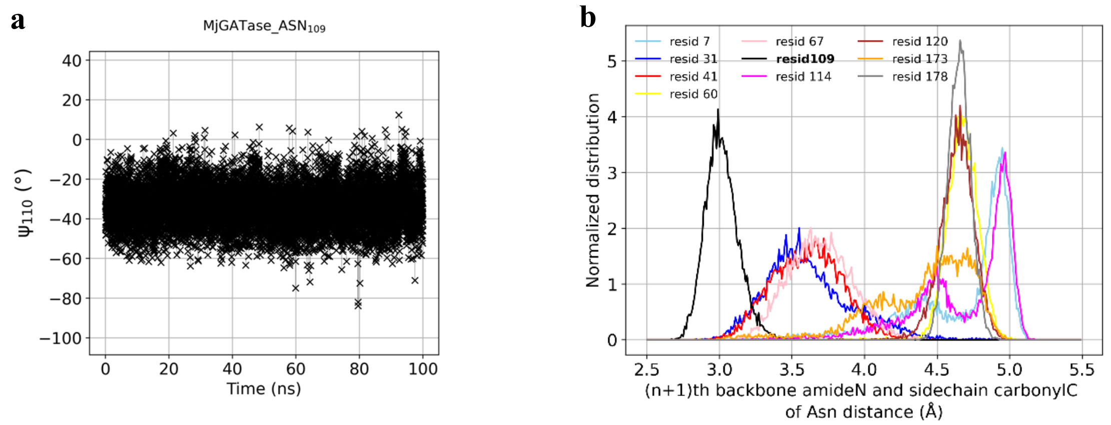


**Supplementary Figure S9. Distribution of D110 ψ values and distance between Asn$_n$ CG and N$_{n+1}$** from classical MD of MjGATase_ASN$_{109}$. (a) Distribution of the backbone dihedral ψ for D110. (b) Normalized distance distributions of the distance between n+1 backbone amide nitrogen and the side chain carbonyl carbon of Asn for all 10 Asn residues in MjGATase. In the inset, 'resid' stands for 'residue'. The residue number is indicated adjacent to resid.

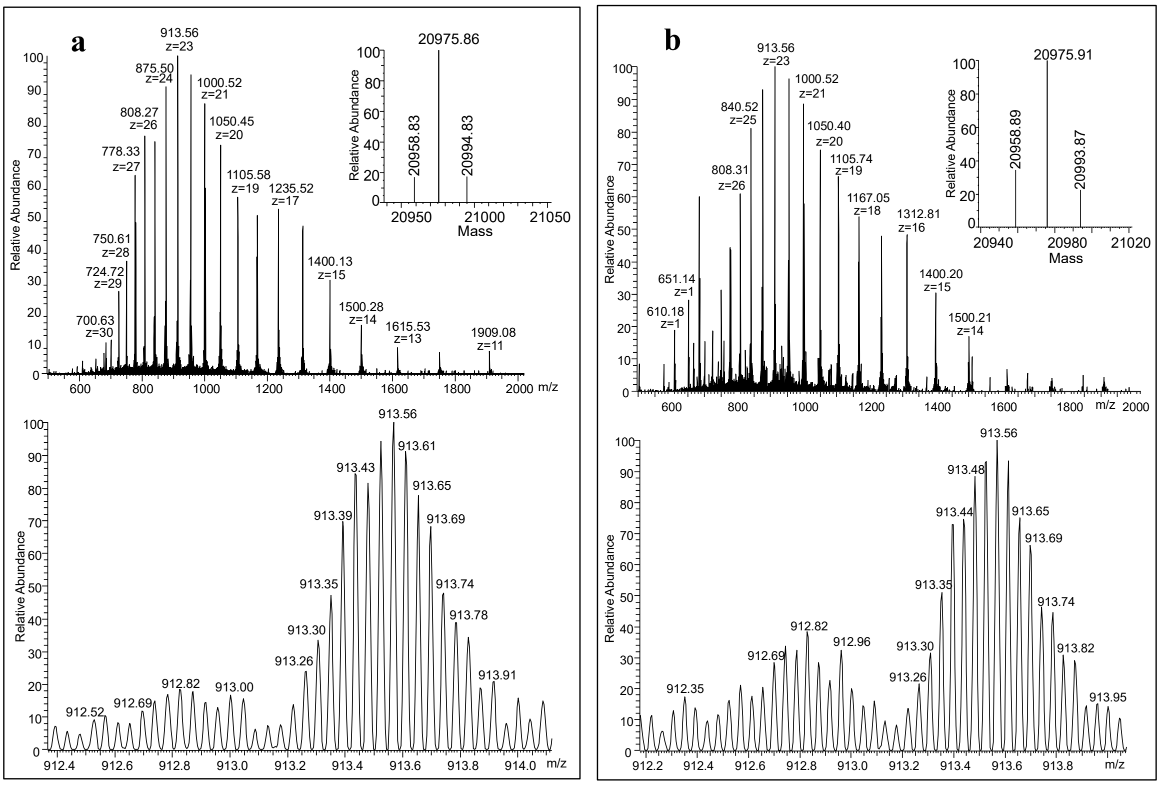


**Supplementary Figure S10. Mass spectra of MjGATase_D110V_K151L ($M_{calc}$ 20975.91 Da) at pH 3, before (a) and after (b) incubation at 60 °C.** (a) Mass spectrum of the protein in pH 3 buffer, 20 °C. Deconvoluted spectrum in the inset shows the monoisotopic mass of a major population with $M_{obs}$ of 20975.86 Da corresponding to the expected mass with Asn109 intact and two minor populations with $M_{obs}$ of 20958.83 Da that is 17 Da lower and corresponding to SNN, and an unassigned $M_{obs}$ of 20994.83 Da corresponding to mass 19 Da higher, from the expected mass. (b) Mass spectrum of the protein in pH 3 buffer after incubation at 60 °C for 16.5 hours. Deconvoluted spectrum in the inset shows the monoisotopic mass of a major population with $M_{obs}$ of 20975.91 Da corresponding to the expected mass with Asn109 intact and two minor populations with $M_{obs}$ of 20958.89 Da that is 17 Da lower and corresponding to SNN, and an unassigned $M_{obs}$ of 20993.87 Da corresponding to mass 18 Da higher from the expected mass. The bottom panel in (a) and (b) shows the expanded spectrum of a single charge state.

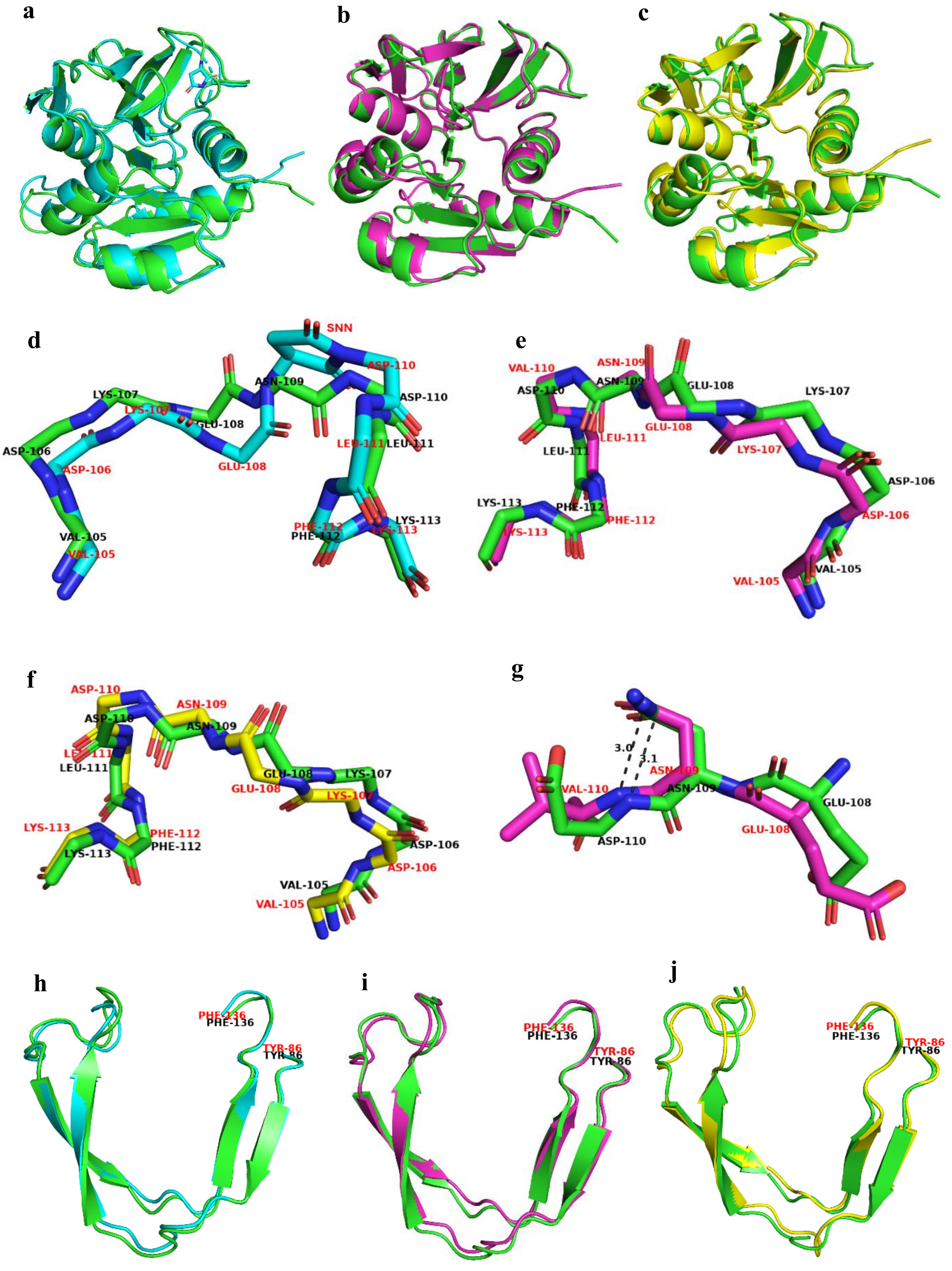


**Supplementary Figure S11. Superposition of the crystal structures of WT and double mutants of MjGATase that have retained Asn109, on to the in-silico generated MjGATase_ASN$_{109}$**. Superposition of the crystal structures of (a) MjGATase_SNN$_{109}$ (cyan), (b) MjGATase_D110V_K151L (purple), and (c) MjGATase_K151L_Y158F (yellow) on the *in silico* generated structure of MjGATase_ASN$_{109}$ (green) that was used for QM/MM studies. Structural superposition of the loop containing SNN/Asn$_{109}$ from residues 105 to 113 in the crystal structures of (d) MjGATase_SNN$_{109}$ (cyan), (e) MjGATase_D110V_K151L (purple), and (f) MjGATase_K151L_Y158F (yellow) on the *in silico* generated structure of MjGATase_ASN$_{109}$ (green). Apart from SNN109, only back bone atoms are shown. (g) Structural superposition of residues 108 to 110 of MjGATase_D110V_K151L (purple) on *in silico* generated MjGATase_ASN$_{109}$ (green). Dashed line denotes the contact between backbone amide of Val/Asp110 and CG of Asn109 with the number indicating distance in Å. Superposition of the β-hairpin segment from residues 86 to 136 in the crystal structures of (h) MjGATase_SNN$_{109}$ (cyan), (i) MjGATase_D110V_K151L (purple), and (j) MjGATase_K151L_Y158F (yellow) on the *in silico* generated structure of MjGATase_ASN$_{109}$ (green). All structure analyses were carried out using PyMOL.

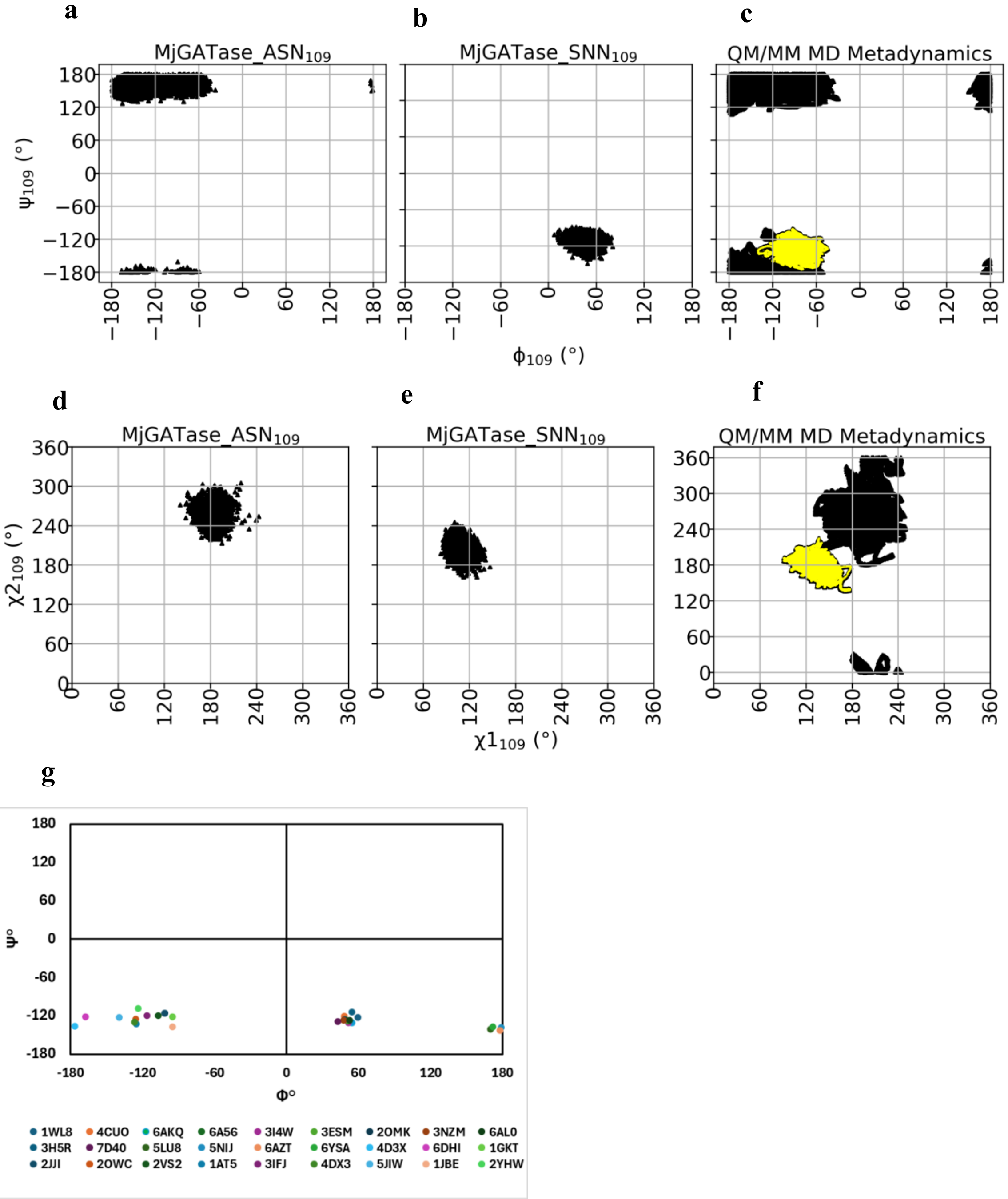


**Supplementary Figure S12. The backbone and side chain dihedral angles of SNN/Asn**. (a) to (c) show the distribution of the Ramachandran angles for the 109$^{th}$ residue in the classical MD (a and b for MjGATase_ASN$_{109}$ and MjGATase_SNN$_{109}$, respectively) and QM/MM MD Metadynamics (c) runs. (d) to (f) show Janin plots for the side chain of the 109$^{th}$ residue in the Classical MD (d and e for MjGATase_ASN$_{109}$ and MjGATase_SNN$_{109}$, respectively) and QM/MM MD Metadynamics (f) runs. In (a) and (d), the 109$^{th}$ residue is Asn, in (b) and (e) it is SNN, and in (c) and (f), the 109$^{th}$ residue changes from Asn to SNN where the SNN containing fractional run length is highlighted in yellow. The product zone, or succinimide containing yellow area, is defined with a distance cutoff (upper bound) of 1.75 Å between the (n+1) backbone amide nitrogen of D110 and the side chain amide carbon of N109. (g) The φ and ψ values of succinimidyl residues in 27 structures in the protein data bank (PDB) are plotted on the Ramachandran map.

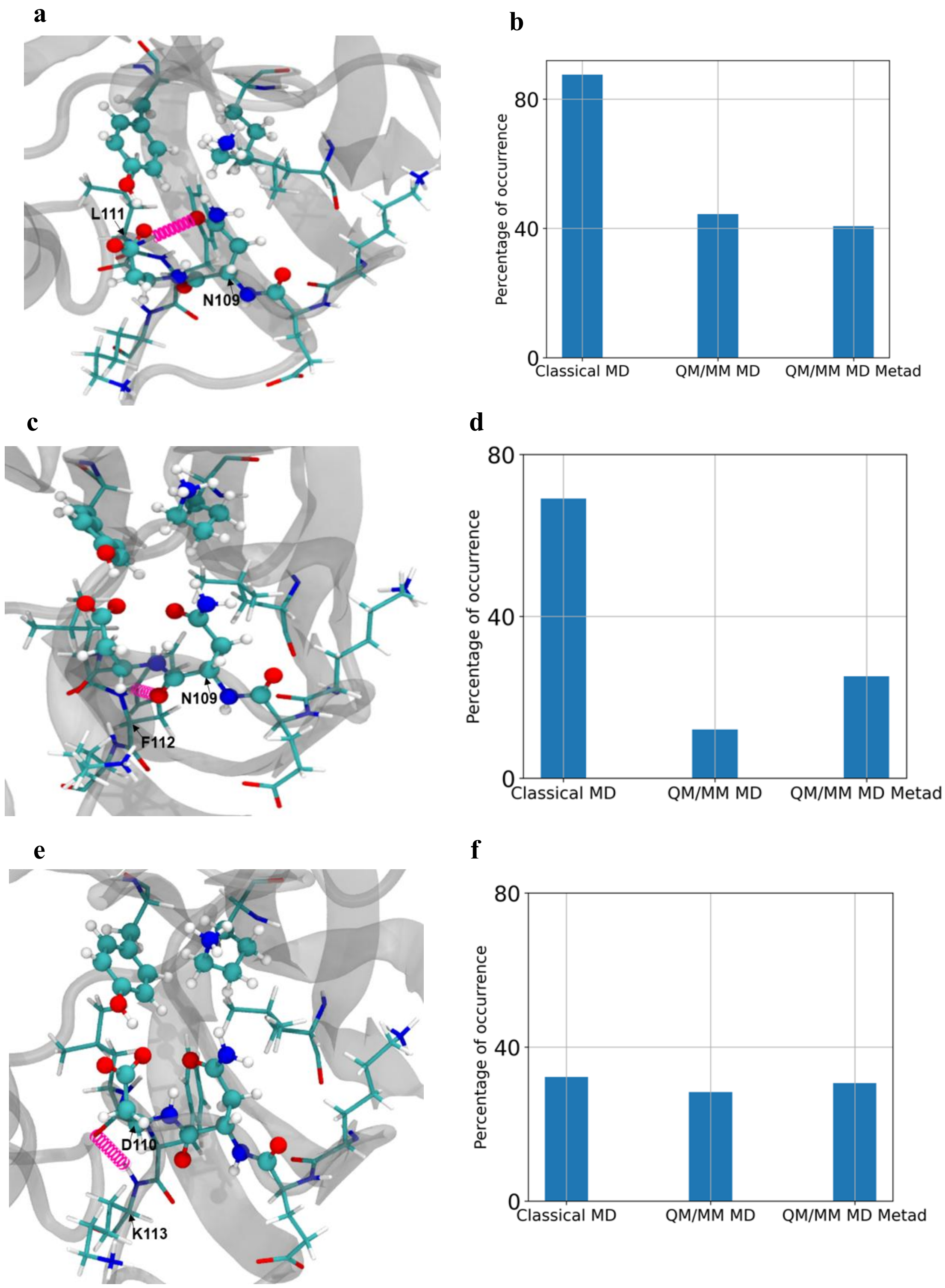


**Supplementary Figure S13. Interactions of Asn/SNN109 and D110 with neighbouring atoms. The interactions shown have maximum frequency of occurrence in the energy minimised MjGATase_ASN$_{109}$**

**structure**. (a) (c) and (e) shows a graphical representation with atoms in QM region in CPK representation (complete residues and some neighbouring residues in Licorice representation). The CA atom of labelled residue is indicated by an arrow. (b) (d) and (f) show the percentage of appearance of the hydrogen bonds in different simulations. Here the y axis shows the fractional count i.e., (count/total_number_of_frames)*100. Classical MD and QM/MM MD contains Asn at the 109$^{th}$ position and for QM/MM MD Metad (QM/MM MD Metadynamics) Asn transforms to SNN in the reaction. (a) and (b) Hydrogen bonding between side chain amide oxygen of Asn/SNN109 and backbone amide H of L111. (c) and (d) Hydrogen bonding between backbone oxygen of Asn/SNN109 and backbone amide H of F112. (e) and (f) Hydrogen bonding between backbone oxygen of D110 and backbone amide H of K113.

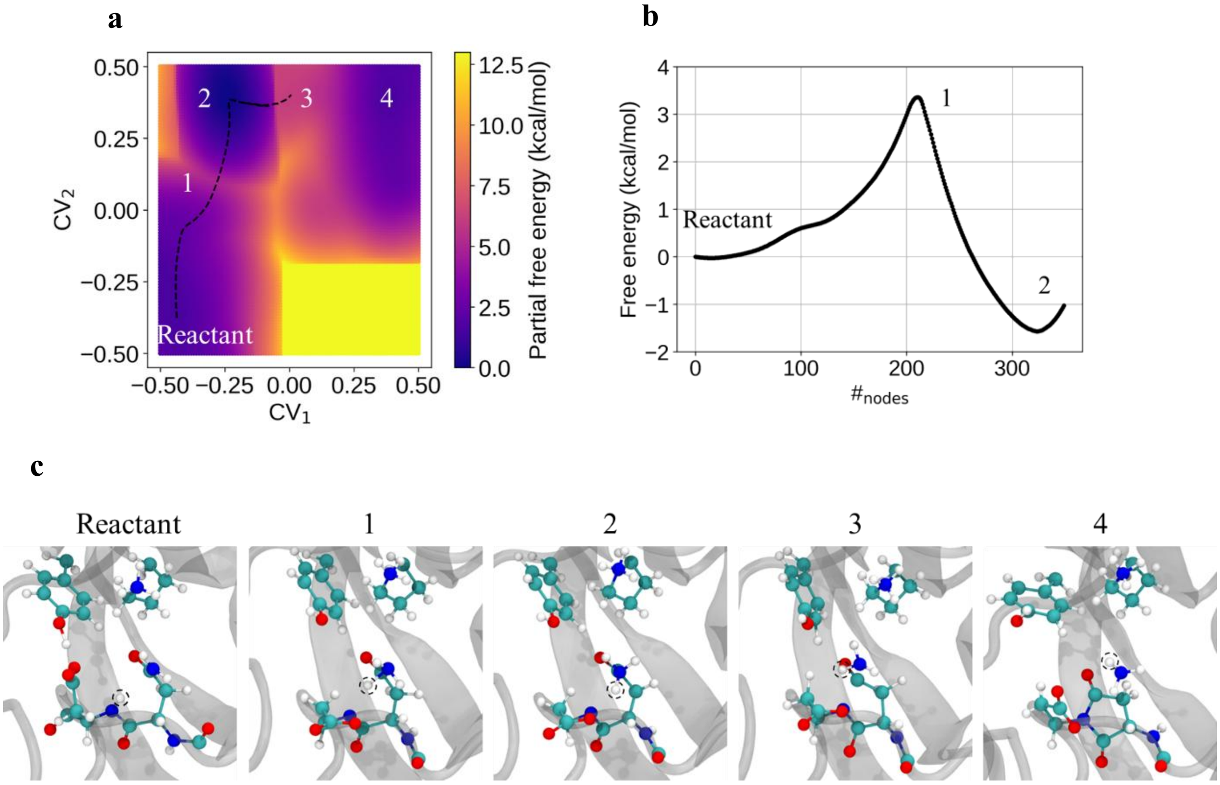


**Supplementary Figure S14. Free energy profile and snapshots of the reaction.** (a) Partial free energy surface for the reaction leading to succinimide formation. For regions having free energy values greater than 13.000364 kcal mol$^{-1}$, the colour used for 13.000364 kcal mol$^{-1}$ has been retained. The minimum free energy path starting from the reactant basin is shown as black dotted line. Four states, from 1 to 4 are highlighted on the surface. (b) The partial free energy profile along the minimum energy path. Reactant state, state 1, and state 2 are indicated. (c) Reactant basin and intermediates in the pathway to succinimide formation. (1) to (4) represent frames from states 1 to 4, respectively from panel (a). The backbone amide proton of D110 is encircled with a black dashed circle.

Table S1. Summary of intact protein mass analysis[a].

| MjGATase protein | Expected monoisotopic mass (Da) | Observed monoisotopic mass (Da) | Mass difference (ppm)[b] | Molecular species[c] | Relative abundance[d] |
|---|---|---|---|---|---|
| Wildtype | 20989.85 | 20989.86 | 0.48 | SNN | 100 |
| | 21006.88* | 21006.83 | 2.38 | Intact Asn | 16.62 |
| | 21021.84 | 21021.87 | 1.43 | SNN+2 methionines oxidised | 15.56 |
| E108L | 20973.89 | 20973.91 | 0.95 | SNN | 100 |
| | 20991.90 | 20991.89 | 0.48 | D/iso D | 18.56 |
|  | 20990.92* | ---- |  | Intact Asn | -- |
| E108Q | 20988.86 | 20988.96 | 4.76 | SNN | 100 |
| | 21020.85 | 21020.95 | 4.75 | SNN+2 methionines oxidised | 17.56 |
| | 21005.89* | 21005.91 | 0.95 | Intact Asn | 17.27 |
| | | 21086.921 | | Unassigned | 15.92 |
| E137L | 20973.89 | 20973.91 | 0.95 | SNN | 100 |
| | 20991.90 | 20991.89 | 0.47 | D/iso D | 22.51 |
|  | 20990.92* | ---- |  | Intact Asn | -- |
| K107L | 20974.84 | 20974.80 | 1.9 | SNN | 100 |
|  | 20991.87* | ---- |  | Intact Asn | -- |
| K151L | 20974.84 | 20974.86 | 0.95 | SNN | 100 |
| | 20991.87* | 20991.84 | 1.43 | Intact Asn | 18.31 |
| Y158F | 20990.89* | 20990.92 | 1.43 | Intact Asn | 100 |
| | 20973.86 | 20973.92 | 2.86 | SNN | 31.67 |
| K113A | 20932.79 | 20932.76 | 1.43 | SNN | 100 |
| | 20949.82* | 20949.73 | 4.29 | Intact Asn | 47.06 |
| D110N | 21006.88 | 21006.89 | 0.48 | D/iso D | 100 |
| | 20988.87 | 20988.88 | 0.48 | SNN | 35.55 |
|  | 21005.90* | ---- |  | Intact Asn | -- |
| D110V | 20990.92* | 20990.93 | 0.48 | Intact Asn | 100 |
| | 20973.89 | 20973.93 | 1.91 | SNN | 91.57 |
| D110P | 20988.91* | 20988.89 | 0.95 | Intact Asn | 100 |
| D110V_K151L | 20975.91* | 20975.91 | 0 | Intact Asn | 100 |
| | | 20959.89 | | Unassigned | 14.7[e] |
| K151L_Y158F | 20975.88* | 20975.84 | 1.91 | Intact Asn | 100 |
| | | 20959.80 | | Unassigned | 11.42[e] |

[a]All studies were carried out using purified protein samples on a Q-exactive HF mass spectrometer. Details of experimental conditions are provided in the methods section.

*Indicates the expected mass of the protein without any modification.

[b]The mass difference is the difference between the observed mass and expected mass and values provided are in parts per million (ppm).

[c]The observed molecular mass has been corelated with the species of MjGATase; *viz.,* with protein having SNN, intact Asn109, hydrolysis product of SNN (D/iso-D), and methionine oxidation.

[d]The most abundant species is 100 with abundance of other species being relative to this value.

[e]While for wildtype and single mutants the abundance cutoff was set to 15%, for D110V_K151L and K151L_Y158F MjGATase mutants the cutoff was set at 10%. Only in the two double mutants even at an abundance cutoff of 10%, a distinct low percentage of species that was above the noise could be observed. In all other cases, lowering abundance cutoff to 10% yielded low abundant species that was at the noise level.

Table S2: Summary of MS/MS analysis of in-gel digested tryptic peptide containing the 109th residue

| MjGATase protein | Observed peptides containing 109th residue | Number of Peptide spectrum matches (PSM) for 109th residue containing | | | Percentage population for 109th residue containing | | |
|---|---|---|---|---|---|---|---|
| | | N | SNN | D/ isoD | N | SNN | D/ isoD |
| Wildtype | VYVDKENDLFK | 9 | 53 | 112 | 2.8 | 34.6 | 62.6 |
| | ENDLFK | | 49 | 30 | | | |
| | VYVDKENDLFKNVPR | | | 13 | | | |
| | KENDLFK | | 6 | 28 | | | |
| | VDKENDLFK | | 2 | 16 | | | |
| K151L | VYVDKENDLFK | 13 | 60 | 131 | 4.9 | 33.8 | 61.3 |
| | ENDLFK | 3 | 45 | 27 | | | |
| | VYVDKENDLFKNVPR | | | 13 | | | |
| | KENDLFK | | 2 | 22 | | | |
| | VDKENDLFK | | 4 | 8 | | | |
| K113A | VYVDKENDLFANVPR | 27 | 100 | 196 | 13 | 31.8 | 55.2 |
| | ENDLFANVPR | 16 | 31 | 8 | | | |
| | AEAEEYALTKVYVDKENDLFANVPR | 2 | | 10 | | | |
| | KENDLFANVPR | 10 | 8 | 24 | | | |
| | YVDKENDLFANVPR | | | 2 | | | |
| | DKENDLFANVPR | | | 2 | | | |
| | VDKENDLFANVPR | 4 | 5 | 8 | | | |
| D110V | VYVDKENVLFK | 57 | 45 | 148 | 26.6 | 20.8 | 52.6 |
| | ENVLFK | 7 | 31 | 19 | | | |
| | VYVDKENVLFKNVPR | 3 | 12 | 4 | | | |
| | AEAEEYALTKVYVDKENVLFK | 6 | | | | | |
| | KENVLFK | 12 | | 13 | | | |
| | VDKENVLFK | 17 | | 18 | | | |
| | YVDKENVLFK | 6 | | 9 | | | |
| | DKENVLFK | 5 | | 12 | | | |
| Y158F | VYVDKENDLFK | 47 | 18 | 43 | 45 | 23.7 | 31.3 |
| | ENDLFK | 15 | 24 | 7 | | | |
| | VYVDKENDLFKNVPR | 6 | | 2 | | | |
| | AEAEEYALTKVYVDKENDLFK | 5 | | | | | |
| | KENDLFK | 7 | 3 | 8 | | | |
| | VDKENDLFK | 9 | 2 | 2 | | | |
| D110N | VYVDKENNLFK | 35 | 11 | 124 | 19.2 | 17.5 | 63.3 |
| | ENNLFK | 5 | 38 | 31 | | | |
| | VYVDKENNLFKNVPR | 7 | | 8 | | | |
| | AEAEEYALTKVYVDKENNLFK | 7 | | 7 | | | |
| | KENNLFK | | | 10 | | | |
| | VDKENNLFK | 2 | 2 | 5 | | | |
| D110P | VYVDKENPLFK | 141 | | | 100 | | |
| | ENPLFK | 33 | | | | | |
| | VYVDKENPLFKNVPR | 3 | | | | | |
| | KENPLFK | 11 | | | | | |
| | VDKENPLFK | 18 | | | | | |
| D110V_ K151L | VYVDKENVLFK | 157 | 5 | 11 | 87.4 | 5.1 | 7.5 |
| | ENVLFK | 25 | 8 | 6 | | | |

| | | | | | | | |
|---|---|---|---|---|---|---|---|
| | VYYVDKENVLFKNVPR | 3 | | | | | |
| | AEAEEYALTKVYYVDKENVLFK | 12 | | | | | |
| | KENVLFK | 7 | | | | | |
| | VDKENVLFK | 18 | | 2 | | | |
| K151L_ Y158F | VYYVDKENDLFK | 85 | 7 | 11 | 82 | 11.2 | 6.8 |
| | ENDLFK | 20 | 11 | | | | |
| | VYYVDKENDLFKNVPR | 9 | | | | | |
| | AEAEEYALTKVYYVDKENDLFK | 6 | | | | | |
| | KENDLFK | 8 | | | | | |
| | VDKENDLFK | 4 | | | | | |

Table S3. List of primers used for the generation of MjGATase mutants. The bases corresponding to the desired mutation are shown in bold letters.

| No. | Primer name | Primer sequence (5' to 3') |
|---|---|---|
| P1 | Y158F_MjGAT_FP | GAAACATAAAACAAAGCCGATT**TTT**GGAGTTCAGTTCCACCCTGAAG |
| P2 | Y158F_MjGAT_RP | CTTCAGGGTGGAACTGAACTCC**AAA**AATCGGCTTTGTTTTATGTTTC |
| P3 | D110V_FP | GGTCTATGTAGATAAAGAAAAC**GTT**TTATTTAAAAACGTTCCAAGAG |
| P4 | D110V_RP | CTCTTGGAACGTTTTTAAATAA**AAC**GTTTTCTTTATCTACATAGACC |
| P5 | D110P_FP | GGTCTATGTAGATAAAGAAAAC**CCG**TTATTTAAAAACGTTCCAAGAG |
| P6 | D110P_RP | CTCTTGGAACGTTTTTAAATAA**CGG**GTTTTCTTTATCTACATAGACC |
| P7 | K151L_MjGAT_FP | GATATATGTCAGGTTGAAGCAATG**CTG**CATAAAACAAAGCCGATTTATG |
| P8 | K151L_MjGAT_RP | CATAAATCGGCTTTGTTTTATG**CAG**CATTGCTTCAACCTGACATATATC |
| P9 | E137L_MjGAT_FP | GTTAAAAAAGTTCCAGAAGGTTTT**CTG**ATTTTAGCTCATTCAGATATATG |
| P10 | E137L_MjGAT_RP | CATATATCTGAATGAGCTAAAAT**CAG**AAAACCTTCTGGAACTTTTTTAAC |
| P11 | K107L_MjGAT_FP | CTAACAAAGGTCTATGTAGAT**CTG**GAAAACGATTTATTTAAAAACG |
| P12 | K107L_MjGAT_RP | CGTTTTTAAATAAATCGTTTTC**CAG**ATCTACATAGACCTTTGTTAG |
| P13 | E108L_MjGAT_FP | CTAACAAAGGTCTATGTAGATAAA**CTG**AACGATTTATTTAAAAACGTTCC |
| P14 | E108L_MjGAT_RP | GGAACGTTTTTAAATAAATCGTT**CAG**TTTATCTACATAGACCTTTGTTAG |
| P15 | T7 FP | TAATACGACTCACTATAGG |
| P16 | T7 terminator RP | GGTTATGCTAGTTATTGCTCAGCG |
| P17# | K151L Y158F_MjGATFP | GATATATGTCAGGTTGAAGCAATG**CTG**CATAAAACAAAGCCGATT**TTT**G |
| P18# | K151L Y158F_MjGATRP | C**AAA**ATCGGCTTTGTTTTATG**CAG**CATTGCTTCAACCTGACATATATC |
| P19 | E108Q_MjGAT_FP | CAAAGGTCTATGTAGATAAA**CAG**AACGATTTATTTAAAAACGTTC |
| P20 | E108Q_MjGAT_FP | GAACGTTTTTAAATAAATCGTT**CTG**TTTATCTACATAGACCTTTG |
| P21 | K113A_MjGAT_FP | GATAAAGAAAACGATTTATTT**GCG**AACGTTCCAAGAGAGTTCAATG |
| P22 | K113A_MjGAT_FP | CATTGAACTCTCTTGGAACGTT**CGC**AAATAAATCGTTTTCTTTATC |

| | | |
|---|---|---|
| P23 | D110N_FP | GGTCTATGTAGATAAAGAAAAC**AAC**TTATTTAAAAACGTTCCAAGAG |
| P24 | D110N_RP | CTCTTGGAACGTTTTTAAATAA**GTT**GTTTTCTTTATCTACATAGACC |

*#Primers P17 and P18 have two codons mutated*

Table S4. Simulation details (CMD, $\#_{Pro-MM}$, $\#_{Pro-QM}$, $\#_{W}$, $\#_{Ions}$, and Prod. stand for Classical Molecular Dynamics, number of protein atoms in MM region, number of protein atoms in QM region, number of water molecules, number of ions, and production run length, respectively. System names, ASN and SNN mean protein structure containing asparagine and succinimide at 109$^{th}$ position, respectively).

| System | Simulation | $\#_{Pro-MM}$ | $\#_{Pro-QM}$ | $\#_{W}$ | $\#_{Ions}$ | Prod. |
|---|---|---|---|---|---|---|
| ASN | CMD | 2973 | - | 11345 | 3 | 100 ns |
| SNN | CMD | 1880 | - | 12777 | 3 | 100 ns |
| ASN | QM/MM MD | 2916 | 61 | 11345 | 3 | 130.245 ps |
| ASN→SNN | QM/MM MD Metadynamics | 2916 | 61 | 11345 | 3 | 56.8995 ps |
